# Macrophage-induced reduction of bacteriophage density limits the efficacy of *in vivo* pulmonary phage therapy

**DOI:** 10.1101/2024.01.16.575879

**Authors:** Sophia Zborowsky, Jérémy Seurat, Quentin Balacheff, Solène Ecomard, Céline Mulet, Chau Nguyen Ngoc Minh, Marie Titécat, Emma Evrard, Rogelio A. Rodriguez-Gonzalez, Jacopo Marchi, Joshua S. Weitz, Laurent Debarbieux

**Author notes:** These authors contributed equally (Sophia Zborowsky, Jérémy Seurat). Correspondence: LD & JSW.

## Abstract

The rise of antimicrobial resistance has led to renewed interest in evaluating phage therapy. In murine models highly effective treatment of acute pneumonia caused by *Pseudomonas aeruginosa* relies on the synergistic antibacterial activity of bacteriophages with neutrophils. Here, we show that depletion of alveolar macrophages (AM) shortens the survival of mice without boosting the *P. aeruginosa* load in the lungs. Unexpectedly, upon bacteriophage treatment, pulmonary levels of *P. aeruginosa* were significantly lower in AM-depleted than in immunocompetent mice. To explore potential mechanisms underlying the benefit of AM-depletion in treated mice, we developed a mathematical model of phage, bacteria, and innate immune system dynamics. Simulations from the model fitted to data suggest that AM reduce bacteriophage density in the lungs. We experimentally confirmed that the *in vivo* decay of bacteriophage is faster in immunocompetent compared to AM-depleted animals and that AM phagocytize therapeutic bacteriophage. These findings demonstrate the involvement of feedback between bacteriophage, bacteria, and the immune system in shaping the outcomes of phage therapy in clinical settings.

## Introduction

Antimicrobials resistance (AMR) is recognized as one of the major threats to global health in the 21^st^ century ^1–3^ with worrisome projections of millions of deaths yearly at a global scale ^4^. Amongst bacterial pathogens, the WHO has identified the need for solutions to treat prioritized threats posed by multidrug resistant (MDR) pathogens that cannot be treated by many commonly used antibiotics ^3,5^.

In response to the AMR crisis, there has been a resurgence of interest in phage therapy, *i.e.* the use of bacterial viruses (bacteriophages or phages) to treat bacterial infections, as an alternative or as a supplement to antibiotics ^6,7^. Pre-clinical studies in animal models have demonstrated the efficacy of phage therapy in the treatment of bacterial infections ^6–8^. While large phase II clinical trials are still required to advance the broad deployment of phage therapy, a steadily increasing number of patients have access to compassionate treatments in both USA and Europe ^9^. A recent retrospective analysis of such 100 patients have shown that their general health improved for more than 70% ^10^.

The killing efficacy of bacteria by phages relies on well-studied mechanisms of adsorption, injection, replication, assembly and lysis whose efficiencies are influenced by the *in vivo* environment (the infected body site). However, the role of the mammalian immune response induced by the pathogen during phage therapy remains uncertain and sometimes contradictory. Several studies performed with murine models of immunosuppression have shown that phage therapy is effective at treating neutropenic mice infected with *Staphylococcus aureus* ^11^ or pathogenic *Escherichia coli* ^12^, but is ineffective for *Pseudomonas aeruginosa*-infected animals ^13,14^. The use of mathematical models has enabled a deeper understanding of these *in vivo* interactions ^15–17^. For example, Roach *et al.* demonstrated that phage therapeutic-induced clearance of *P. aeruginosa* from the lungs requires synergistic action of phages with neutrophils ^14^, via a mechanism termed immunophage synergy. Another study combining data and mathematical modeling showed the efficacy of the immunophage synergy in treating *E. coli* pulmonary infections was dependent on the size of the inoculum ^18^. The prominent role of neutrophils in the lungs contrast with their density as they represent about 2% of immune cells in this organ ^19^, in contrast to alveolar macrophages (AM), which account for 55% of immune cells in the lungs of healthy humans ^20^.

AM are the first line of cell-mediated defenses against microorganisms inhaled into the respiratory system as resident phagocytes ^21,22^. AM are also the predominant immune cells present in the alveoli in homeostasis ^22,23^ and have several critical functions during infection, participating in clearance of pathogens and repair of the host tissue damages ^24,25^. *P. aeruginosa* pneumonia initiates with the activation of AM through recognition of the LPS by TLR-4 and the flagellin by TLR-5, followed by the secretion of chemokines and cytokines that recruit neutrophils ^26–28^. Nevertheless, studies using AM-depleted animals (different mouse strains and depletion levels) showed mixed outcomes leaving uncertainties about the role of AM in managing *P. aeruginosa* ^29–31^. Furthermore, the direct interactions between AM and phages and their effect on phage therapy are poorly understood, particularly given that the innate immune system could have negative interactions with phage particles that can be phagocytized by neutrophils, monocytes, tissue macrophages and dendritic cells ^32^. However, the response of phagocytizing cells to phage particles is not ubiquitous and some phages do not seem to elicit any interaction with macrophages ^32,33^.

Here, we sought to assess the role and interactions of AM in the clearance of *P. aeruginosa* during acute pneumonia and its treatment by phage. We combined an *in vivo* and mathematical modeling approach to conceptualize and experimentally test the interactions between pathogenic bacteria, therapeutic phage and the innate immune system (including macrophages and neutrophils). As we show, although AM depletion negatively impacts survival of mice given exposure to *P. aeruginosa* in the absence of phage therapy, the depletion of AM can have unexpected benefits to the efficacy of phage infection and lysis leading to reduced levels of *P. aeruginosa* in phage-treated animal hosts.

## Results

### Infection in alveolar macrophage-depleted mice

In previous studies, the virulent phage PAK_P1 (93 kb, *Pakpunavirus*) has been demonstrated to efficiently treat *P. aeruginosa* pneumonia caused by strain PAKlumi (allowing *in vivo* recording of bioluminescence) ^14,34^. With the same phage and bacterium system we aimed to investigate the role of AM using an *in vivo* depletion approach relying on the intra-tracheal administration of clodronate in liposomes at a concentration known to cause macrophage apoptosis ^21,35^. We found that 96 h after a single administration of clodronate the AM population decreases in median (min-max) by 98% (95%-99%), while we did not observe an additional significant effect on lung nor blood neutrophils and dendritic cells (DCs) (Fig. 1A, Sup. Fig. 1 and 2).

**Figure 1.**
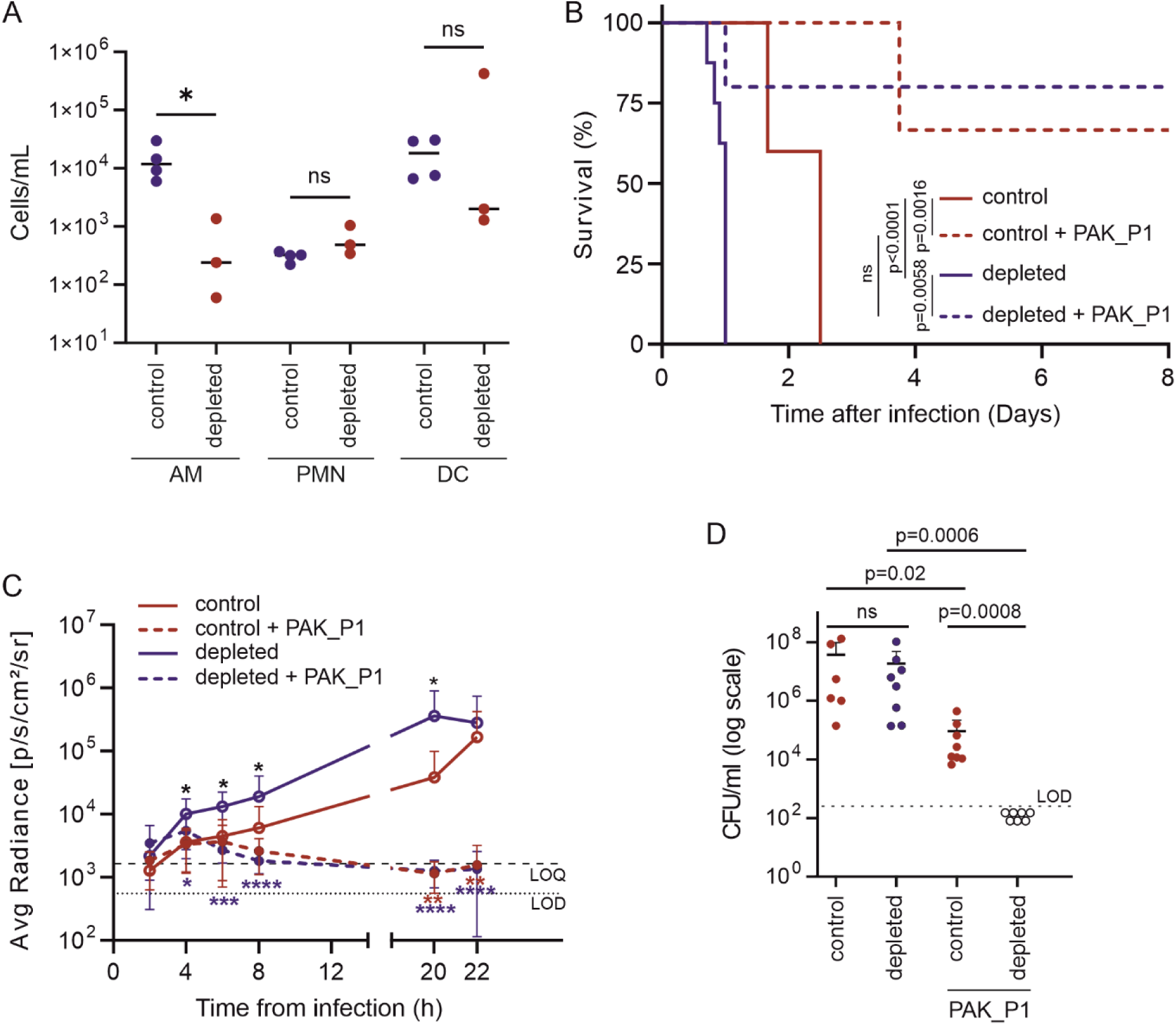
Phage therapy in the macrophage depleted host. (A) Flow cytometry analysis of bronco alveolar lavages (BAL) from mice that were given 96 h before clodronate liposomes (depleted) or empty liposomes (control) by intra-tracheal instillation (n=4 each). Cell debris and doublets were first excluded based on CD45 staining. Among the CD45+ population Ly6G+ was used to gate PMN, in the Ly6G-population, the CD11b-correspond to dendritic cells (DC) and alveoli macrophage (AM). The DCs were gated as F4/80- and CD11c+ low while the AM as F4/80+ and CD11c+ high. AM were significantly depleted (p=0.043), whereas PMN and DC were not affected (Mann-Whitney U test). (B) Survival of acute respiratory infection caused by *P. aeruginosa* (5 x 10^6^ CFU of strain PAKlumi) of AM-depleted mice (clodronate, n=8) and control mice (n=5), p <0.0001 Mantel-Cox log rank test) in absence of phage treatment (full lines). With phage PAK_P1 (5 x 10^7^ PFU) administered intranasally at 2 h p.i. (dashed lines) in AM-depleted (n=5) and control (EL) (n=6) mice, phage treatment significantly improved their survival (p =0.0058 and 0.0016, Mantel-Cox log rank test, for AM-depleted and control mice, respectively). Data were assembled from 5 independent experiments. (C) The colonization kinetics of the bioluminescent strain PAKlumi in the lungs is plotted as mean radiance (p/s/cm^2^/sr) over time. Groups of mice were infected and treated as in panel B with control (n=6), AM-depleted (n=8), phage-treated control (n=9) and phage treated AM-depleted (n=13). Statistical analysis displays unpaired group comparison (Mann-Whitney U test), with stars indicating: in black comparison of untreated groups between depleted and control, in red comparison of control groups between untreated and phage-treated and in blue comparison of AM-depleted groups between untreated and phage-treated. *p<0.05, **p<0.01, ***p<0.005, ****p<0.001. LOQ, limit of quantification. LOD, limit of detection. Data were assembled from 6 independent experiments. (D) Bacterial counts (CFUs in log scale) from lung homogenates taken from mice (same animals as in panel C) 22 h post infection. LOD, limit of detection (p values (Mann-Whitney U test) for between groups comparisons are displayed; ns, not significant). Data were assembled from 5 independent experiments.

We then compared the virulence of the *P. aeruginosa* strain PAKlumi in AM-depleted mice with control mice that received empty liposomes (EL). We found that 100% of AM-depleted mice had reached the compassionate end point within 24 h post-infection (p.i.), while it took over 48 h for the EL group (Fig. 1B). The dynamics of progression of the infection in the lungs was obtained through bioluminescent records over time for all animals showing that the average radiance between the two groups was statistically different from 4 h to 20 h p.i. (Fig. 1C). However, *in vivo P. aeruginosa* growth rate comparisons based on the luminescence recordings between the EL and AM-depleted groups showed no significant differences. Furthermore, lungs were recovered at compassionate end points to measure *P. aeruginosa* densities by colony forming units (CFUs) (Fig. 1D). We observed no significant difference in the CFUs levels between the two groups as suggested by the bioluminescent records at 22 h p.i. before sacrifice. Overall, AM depletion did not induce a significant effect on the *in vivo* growth rate of *P. aeruginosa* but AM depletion did have a significant and negative impact on survival. These results point towards the involvement of AM, as well as other immune cells including neutrophils, in the control and elimination of *P. aeruginosa*.

### Phage therapy rescues macrophage-depleted mice infected by *P. aeruginosa*

In order to assess the involvement of AM in the efficacy of phage treatment we administered a single dose of phage PAK_P1 at a multiplicity of infection (MOI) of 10 (related to the inoculum of strain PAKlumi) 2 h p.i., in both AM-depleted and EL mice. In EL mice the phage treatment increased the survival from 0% at 2.5 days to 67% at 8 days p.i., while in AM-depleted mice survival jumped from 0% at 24 h to 80% at 8 days p.i. (Fig. 1B). Therefore, the phage treatment increased significantly the survival in both AM-depleted (p=0.0058) and EL mice (p=0.0016) compared to the untreated groups. The survival rate at 8 days was not significantly different between the two phage-treated groups suggesting a similar efficacy of the antibacterial action of the phage. The bioluminescent recordings over time confirmed the rapid action of the phage as average bioluminescence values reached nearly the limit of quantification (LOQ) of 2000 p/s/cm²/sr equivalent to 5 x 10^5^ CFU/mL at 8 h p.i. (ie. 6 h post phage administration) (Fig. 1C) and were below the LOQ at 22 h p.i. in both phage-treated groups. Unexpectedly, the CFUs obtained from lungs of phage-treated animals sacrificed at 22 h p.i. showed that in the AM-depleted group no *P. aeruginosa* could be detected (limit of detection of 2.5 x 10^2^ CFU/mL) in sharp contrast with the EL group that had levels around 10^5^ CFU/mL (Fig. 1D). This large difference in CFUs, although not detected from the luminescence data (values below the LOQ), indicates that the depletion of AM increases the clearance of bacteria by phage and suggests that AM may negatively impact phage therapy efficacy.

### Mathematical model of phage-immune feedback during phage therapy recapitulates *in vivo* dynamics in immunocompetent and AM-depleted mice

Building upon an immunophage synergy model ^14^ of *in vivo* phage therapy, we developed a mathematical model that includes the growth of bacteria, phage infection, lysis of susceptible bacteria, as well as the recruitment and clearance of bacteria by immune effector cells (neutrophils), which are themselves recruited by AM (Fig. 2 and Sup. methods for model specification including equations and parameterization). The parameterization of the *in vivo* phage therapy model was enabled by leveraging data obtained from immunocompetent mice (n=44) infected by strain PAKlumi (inoculum of 10^7^ CFU). Bioluminescence recordings were compared to equivalent time-point CFU measurements from plating to establish a correlation of luminescence with CFU data (adj-R²=0.71, Sup. Fig. 3A), which was further used to convert *in vivo* measurements for direct model-data comparison. Given the richness of these longitudinal data (Sup. Fig. 3B), as well as individual phage loads (Sup. Fig. 3C), immune cells data (Fig. 1A) and the observations previously described in the subsections and features of the immunophage synergy model ^14^, we set out to identify parameters that could recapitulate treatment dynamics in multiple conditions. Using normalized prediction distribution errors ^36^ from state-of-the-art practice in mixed effect modeling (Sup. Fig. 4), we find that a common set of parameters in our *in vivo* phage therapy model can summarize observed time series dynamics in EL and AM-depleted mice both without or with phage while accounting for between-mouse variability (Fig. 3) and longitudinal data from all infected mice (Sup. Fig. 5). We interpret model-data fits to mean that our baseline model can jointly explain the observed dynamics of bacteria, neutrophils, and AM in multiple conditions, insofar as phage rapidly proliferate in treated animals, even if there remains uncertainty in the extent to which there may be direct, negative interactions between components of the innate immune system and phage.

**Figure 2.**
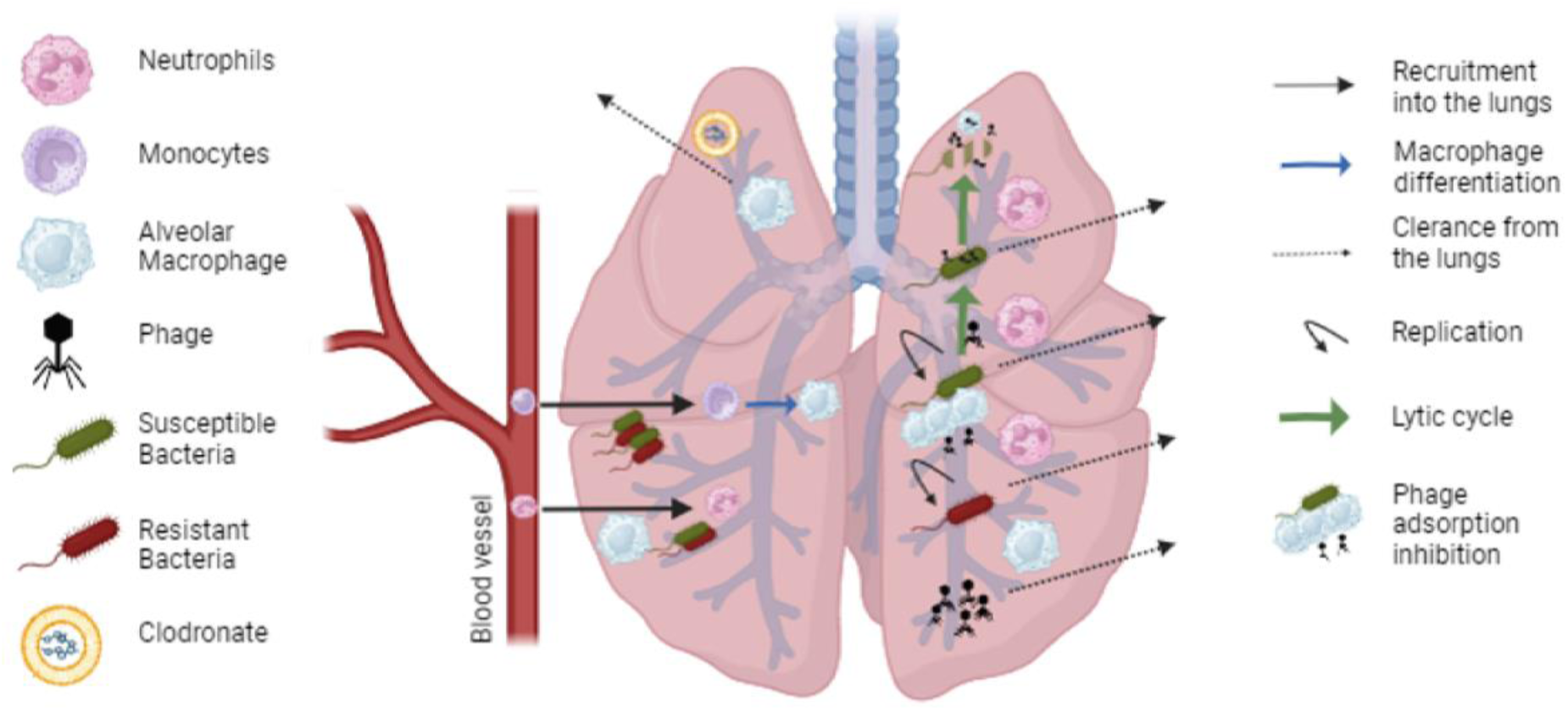
Schematic model of bacteria-phage-macrophage-neutrophils interaction in the lungs. An immune cell close to the arrow represents its impact on the corresponding process. Created in BioRender.com. A full list of equations and parameterization are found in Sup. Methods.

**Figure 3.**
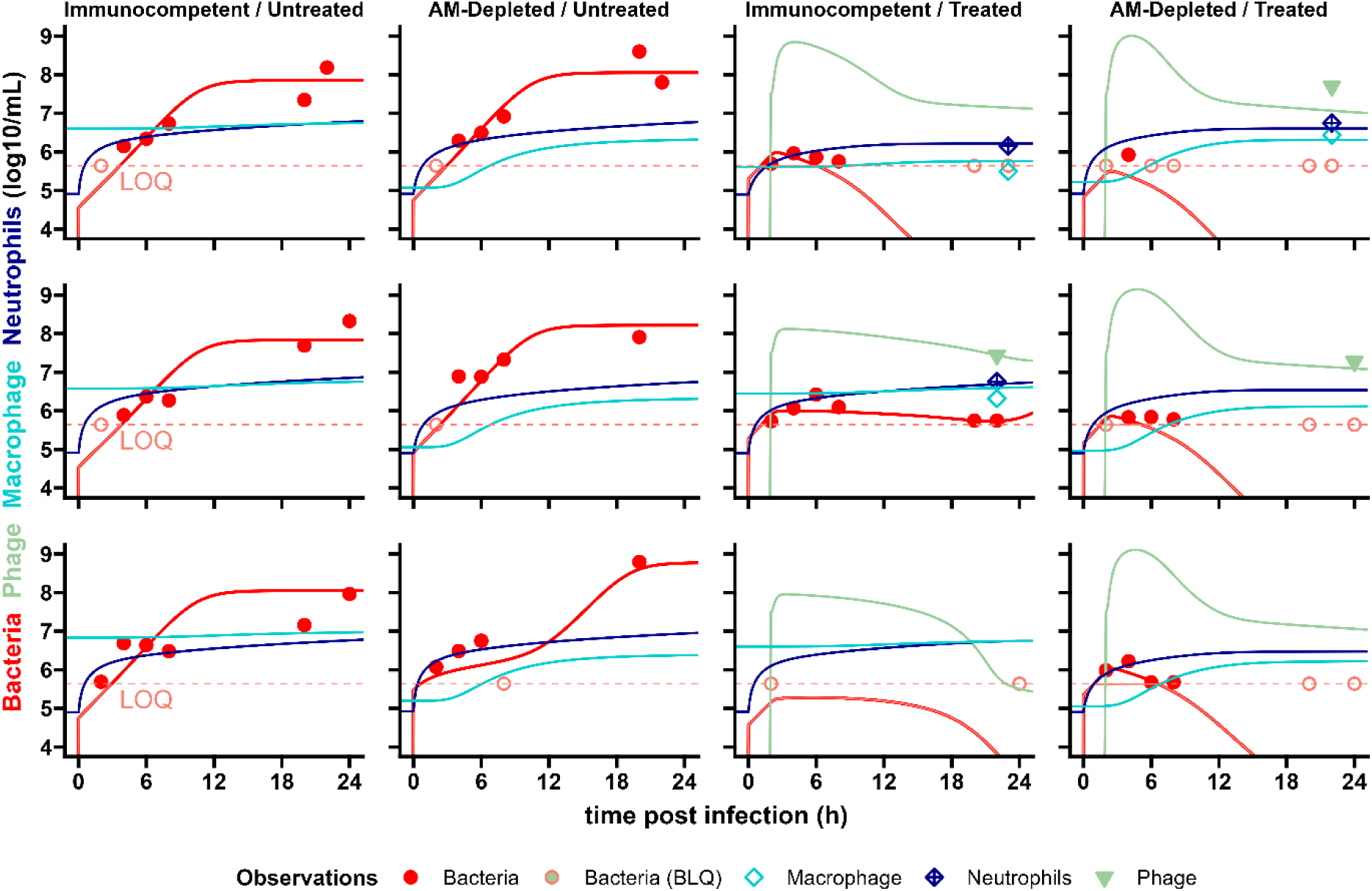
Model of bacteria-phage-macrophage-neutrophils interaction in the lungs. Individual model predictions and data for 3 mice from each group ranked in columns with corresponding headers. Bacteria are in red, phage in green, alveolar macrophage (AM) in light blue, and lung neutrophils in dark blue. Lines are model-based individual dynamics, dots are bacterial densities, triangles are phage concentrations, crossed squares are lung neutrophils and squares are AM. The light red horizontal dashed lines represent the limit of quantification (LOQ) for bacterial density (LOQ = 5.6 log_10_ CFUeq/mL), light red circles crossed by this line at their center are below the limit of quantification (BLQ) of bacterial density and light red lines are BLQ of bacterial load model-based predictions (i.e. the discrepancy between data and model cannot be evaluated from this plot in case of BLQ data).

### Macrophage reduction of phage densities is essential to model *in vivo* data fits

To incorporate the faster progression of the infection observed in AM-depleted compared to EL mice, we tested two, non-exclusive hypotheses by varying the parameters of these processes: (i) phagocytosis of *P. aeruginosa* is eliminated in AM-depleted relative to EL mice; (ii) neutrophil recruitment is reduced in AM-depleted relative to EL mice (Fig. 2, Sup. Methods: Eq. S1-S15). According to the model the difference in infection dynamics are better explained by a delay in recruitment of neutrophils rather than by reduced phagocytosis of bacteria in AM-depleted mice. The best model included a reduction of 50 % of the neutrophil recruitment rate in absence of AM. When adding the component of phagocytosis of *P. aeruginosa* by macrophages the model predictions did not align better with the observed infection dynamics than without phagocytosis (corrected Bayesian information criterion of 3359 vs. 3358, respectively); therefore, we removed AM phagocytosis of *P. aeruginosa* from the model. We incorporated the killing activity of the phage in our model to identify putative mechanisms to explain how phage therapy efficacy increased, albeit more so in AM-depleted than in EL mice. In the model framework, phage rapidly absorb and lyse bacteria. Bacterial growth is then controlled by a combination of phage and neutrophils that are progressively recruited in the lungs.

We next hypothesized that the reduced clearance of *P. aeruginosa* in the lungs of EL mice was caused by the clearance of phage mediated by AM – if so, this would imply that fewer phage were available to infect and lyse *P. aeruginosa*, as observed 22 h p.i (Sup. Fig. 3C). We used the mathematical model to estimate the AM-mediated decay rate of phages, assuming no other interaction between phage and macrophage. We compared CFU densities in the lung to predicted median bacterial dynamics from a mathematical model incorporating phage kinetics in an *in vivo* context. This model included dynamics of pulmonary bacteria, phage, AM and neutrophils while assuming various AM-mediated decay rates. We found that a decay rate of phage PAK_P1 around 8 times higher in EL compared to AM-depleted animals would explain the difference in the CFU data between EL (Fig. 4A) and AM-depleted (Fig. 4B) mice. Indeed, in EL mice every lower decay rate resulted in predicting lower bacterial loads compared to observed data, whereas a higher decay rate led to a late emergence of resistant bacteria, which is not compatible with the radiance data (Fig. 1B) and survival (Fig. 1C). In AM-depleted mice, due to the rapid adsorption and lytic parameters of the phage, all the AM-mediated decay rate values between 1 and 16 times the natural decay (i.e. not AM-mediated decay) rate were compatible with the non-detectable bacteria at 22 h p.i (Fig. 4B). Hence, the mathematical model suggests that AM could reduce phage densities *in vivo* which would have deleterious effects on the efficacy of phage therapy in immunocompetent murine hosts but, in contrast, synergistic effects in AM-depleted murine hosts.

**Figure 4.**
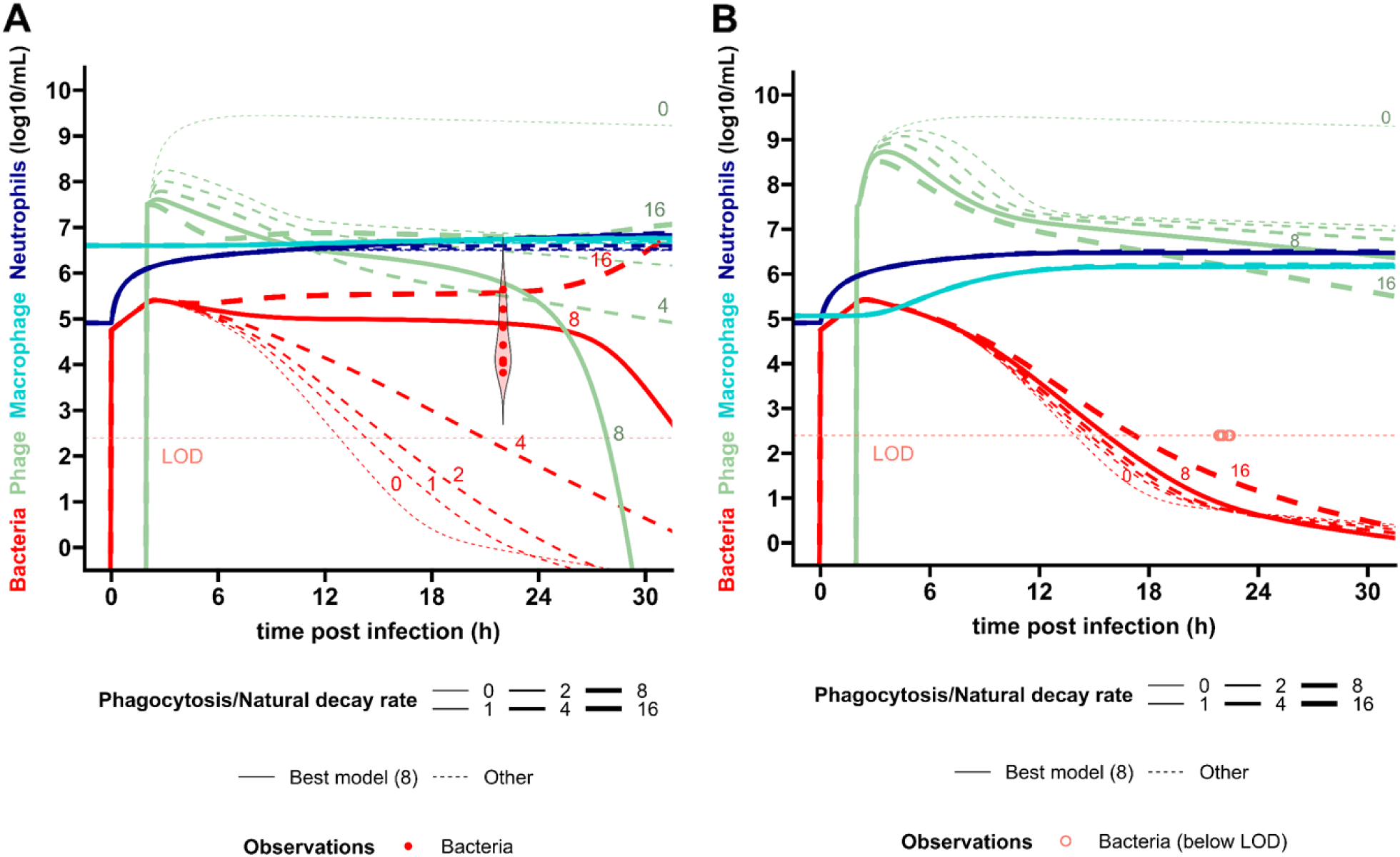
Median dynamics with various AM-mediated phage decay rate parameter values after infection and phage treatment. Dynamics of bacteria (red), phage (green), alveolar macrophage (AM, light blue), and lung neutrophils (dark blue) based on the model and its median parameters (Fig. 2A, Sup. Methods), assuming no interaction between macrophages and phage except AM-mediated decay. Each panel corresponds to the different mice groups according to their immune status: (A) immunocompetent control (*i.e.*, empty liposome), (B) AM-depleted (clodronate). Line widths increase depending on the AM-mediated decay rate (multiplied by the macrophage density): 0,1,2,4,8 and 16 times the phage decay in absence of AM, from the thinnest to the widest line. This number is also indicated above the corresponding median bacteria and phage dynamics. The AM-mediated/natural decay rate value leading to the best model (*i.e.*, 8) is represented by solid lines, while the others are represented by dotted lines. The violin and the dots represent the distribution and values of observed CFUs. The limit of detection (LOD) for CFUs is represented by the light red horizontal dashed line (LOD = 2.4 log_10_ CFU/mL). For AM-depletion, all the observations were below the LOD.

### Phagocytosis is involved in phage decay

In order to test if AM reduced phage concentration in the lungs, we administered a single dose of phage PAK_P1 to AM-depleted and EL mice and sacrificed animals over time to assess phage decay rates *in vivo*. We found that the levels of PFUs were significantly lower in the EL compared to the AM-depleted mice across all time points (Fig. 5A). Performing nonlinear regressions, the phage decay rate was significantly 3 times lower (3.1, CI_95_: [2.9, 3.2], p<0.0001) in the depleted compared to the EL groups (0.023 [0.016,0.029] h^-1^ vs. 0.069 [0.067, 0.071] h^-1^, which corresponds to elimination half-lives of 30 h and 10 h, respectively. Moving one step further we quantified by qPCR the amount of phage DNA recovered from AM sorted (CD45, CD11c and Singlec-F positive cells) from bronco-alveolar lavages (BAL) of lungs of mice exposed to either PBS or PAK_P1. The five mice that were administered intranasally phage PAK_P1 had a mean number of phage DNA copies per AM ranging from 2.6 to 6.0 while no phage DNA was detected in PBS-treated mice (Fig. 5B). Altogether, these data suggest that phage are being removed from the lungs by AM at a rate which is consistent with a previous study evaluating phage decay in immunocompetent hosts (0.07 h^-1^)^14^.

**Figure 5.**
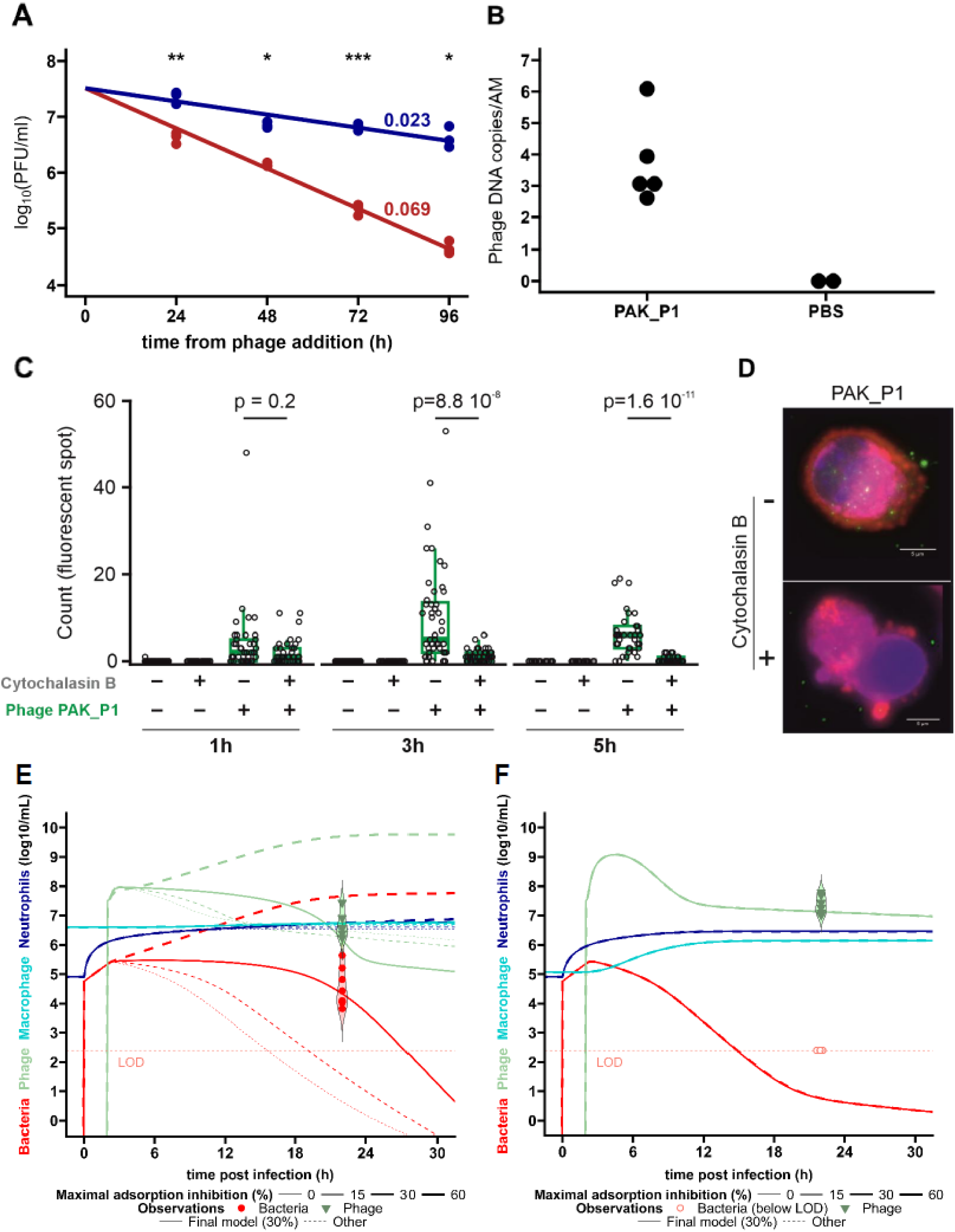
AM-associated phage decay and adsorption inhibition assumption. (A) Uninfected mice were given clodronate (blue) or empty liposomes (red) by intra-tracheal instillation (n=3-4 each) 4 days prior giving phage PAK_P1 (5 x 10^7^ PFU) at t = 0 h. Lungs were collected at 24, 48, 72 and 96 h and homogenized before plating to determine PFU. Data fitted using a monoexponential decay model in the two different groups, with the colored value indicating the corresponding decay rate above each line. *: <0.05, **: <0.01, ***: <0.001, (Mann-Whitney U test). (B) Uninfected mice were given intranasally PBS (n=2) or phage PAK_P1 (5 x 10^7^ PFU) at t = 0 h (n=6) and broncho alveolar lavage was performed 4 h post-administration to sort and count AM and quantify the amount of phage DNA by qPCR. (C) Counts of phage-associated fluorescent signals in individual AM collected from uninfected mice and incubated without (control) or with phage PAK_P1 in absence or presence of cytochalasin B at indicated time points (n=13 to 47 for each condition). (D) Representative images from AM incubated with phage PAK_P1 in presence or absence of cytochalasin B. (E,F) Median dynamics with various adsorption inhibition parameter values after infection and phage treatment for (E) immunocompetent control (i.e. empty liposome) and (F) AM-depleted mice. Dynamics of bacteria (red), phage (green), alveolar macrophage (AM, light blue), and lung neutrophils (dark blue) based on the model and its median parameters (Fig. 2A, Sup. Methods), assuming phagocytosis rate value from the inference of the data of Fig. 4A. Line widths increase depending on the maximal adsorption inhibition by AM: 0%, 15%, 30% and 60%, from the thinnest to the widest line. The maximal adsorption inhibition value leading to the best model (*i.e.*,30%) is represented by solid lines, while the others are represented by dotted lines. The violin and dots represent the distribution and values of observations: CFUs in red and PFUs in green. The limit of detection (LOD) for CFUs is represented by the light red horizontal dashed line (LOD = 2.4 log_10_ CFU/mL) For AM-depletion, all the observations were below the LOD.

Next, we performed *in vitro* assays by first incubating phage PAK_P1 with murine macrophage J774 cell line at three phage:cell ratios during 24 h. We observed no significant differences in phage titers (Sup. Fig. 6A). Second, we incubated phage PAK_P1 with AM collected from uninfected mice in the absence and presence of cytochalasin B, a drug known to inhibit phagocytosis. We observed a significant increase of phage-associated fluorescent signals in AM in the absence of cytochalasin B compared to its presence at times 3 h and 5 h post-incubation demonstrating a significant phagocytosis of phages by AM (Fig. 5C,D, Sup. Fig. 6B,C). These data suggest that either macrophage J774 cell line are unable to phagocytose phage in contrast to AM or, more likely, that phagocytosis of phages was not detectable by direct plating.

However, the mathematical model predicts that the accelerated decay rate of phage is not high enough to explain the 3-log difference in CFUs observed at 22 h p.i. between these EL and AM-depleted mice (Fig. 1D), suggesting the presence of additional inhibitory mechanisms beyond phagocytosis. Macrophages have been shown to “cloak” areas of injury or infection in order to reduce neutrophils recruitment and sequential neutrophil-mediated inflammatory damages ^37,38^. Macrophage that block access to bacteria might also reduce contact by phage, reducing adsorption. By setting values for natural phage decline and AM-mediated decay rate to the observed values, we investigated a range of AM-induced reduction of phage adsorption within our mathematical model. In EL mice, the combination of the removal of phages at a rate of 0.069 h^-1^ with a reduction of phage adsorption by 30% generated *P. aeruginosa* clearance rates corresponding to the experimental CFUs in this group whereas a lower value such as 15% led to faster clearance than observed and a higher value such as 60% led to bacterial control (Fig. 5E). In AM-depleted, the different adsorption inhibition values, including 30%, all recapitulate the bacterial clearance at 22 h p.i., since the low density of AM was insufficient to inhibit phage adsorption in the first 10 h p.i., allowing phages to multiply and hence rapidly clear bacteria (Fig. 5F). Interestingly, the combination of the two AM-phage processes (phage decay and adsorption inhibition) could explain the small difference (about 1 log) of phage density between EL and AM-depleted mice (Sup. Fig. 3C).

Overall, the negative effects of AM on phage efficacy result in the median bacterial load stabilizing during the 18 h following phage administration in the EL mice (Fig. 5E) versus 4 h in the AM-depleted animals (Fig. 5F). This negative effect of AM on phage concentration is counterbalanced by the infection parameters of phage PAK_P1 (rapid adsorption rate of 1.2 10^-7^ mL/PFU/h, lytic cycle of 20 minutes and burst size of 100 ^14^), leading to decreasing bacterial load. From our model-based simulations accounting for inter-individual variability, differences were noted between the EL and the AM-depleted phage-treated groups (Fig. 6). Indeed, in the EL group, our model predicts a bacterial rebound of over 10^7^/mL for at least 25% of individuals 30 h post-infection, whereas in the AM-depleted group, over 75% of individuals had bacterial load below 10^4^/mL. Notably, the antagonistic interactions between AM and phage could result in loss of bacterial infection control, depending on the phage dose. Our model predicts a possible rebound in bacterial load at MOI 0.1, i.e., with a dose 100 times lower than that used in our study. For AM-depleted individuals, the decline in bacterial load becomes slower as the phage dose decreases, without showing a rebound even at MOI 0.01 (Sup. Fig. 7).

**Figure 6.**
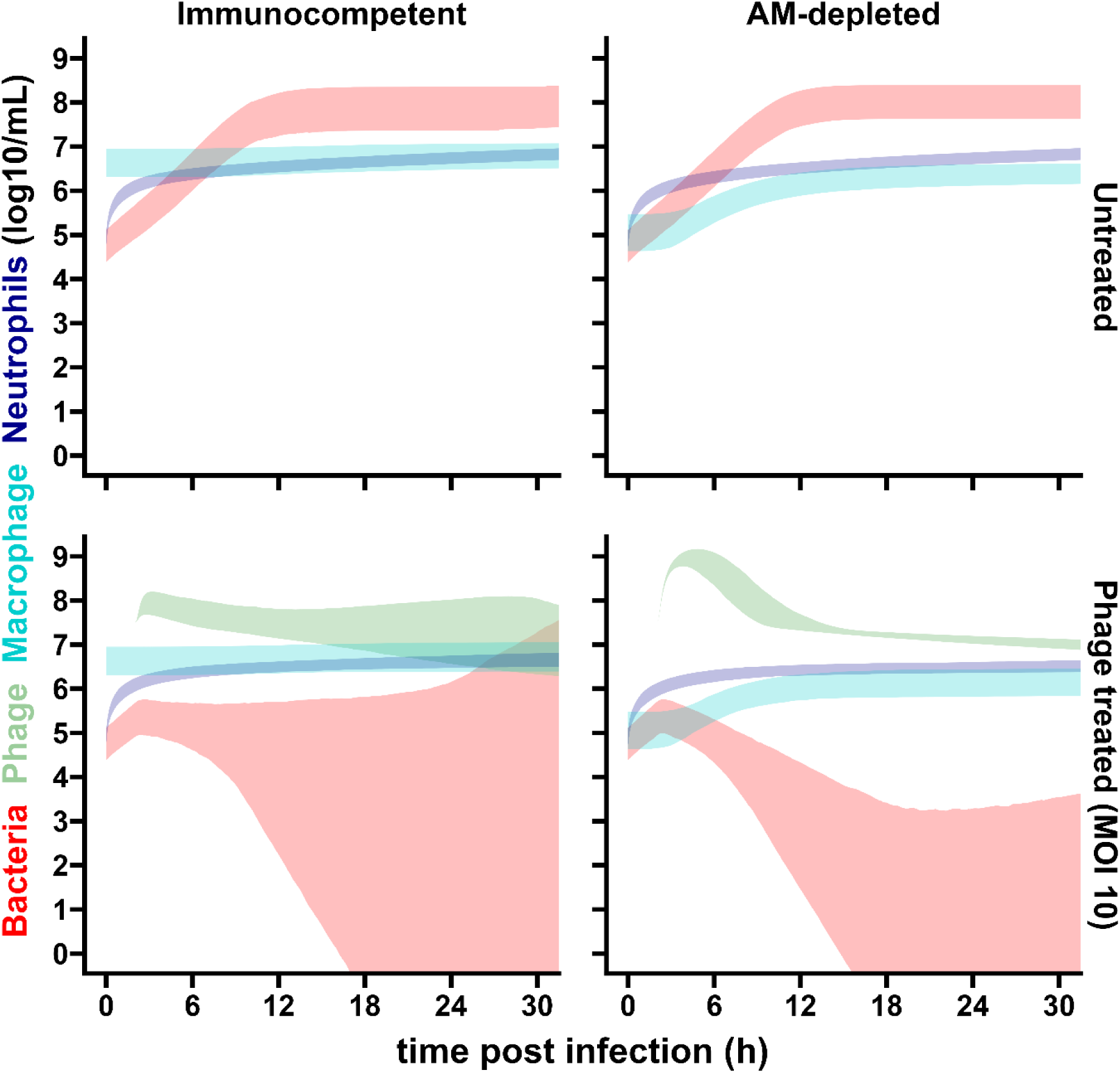
Inter-individual variability in the dynamics of bacteria, phage, alveolar macrophage and lung neutrophils. Based on the model (Fig. 2) and its median and variability parameters (Sup. Methods), simulations for 1000 individuals in each group are shown by areas covering interquartile interval (Q1-Q3). Each panel corresponds to the different mice groups according to their immune status and phage treatment.

## Discussion

In this study we examined the role of alveolar macrophage (AM) during *P. aeruginosa* infection and treatment with phage leading to two central conclusions. First, based on survival analysis we infer that AM are a necessary immune component for survival of mice to *P. aeruginosa* infection. Even though the initial progression of *P. aeruginosa* infection is faster in the absence of AM – a feature which can be explained by the delayed recruitment of AM-induced neutrophils into the lungs – the bacterial load is not significantly different at the compassionate end point. Thus, we concluded that the reduced survival in the absence of macrophages is not related to the lack of phagocytosis of *P. aeruginosa* by AM. Second, we found that phage therapy was curative in both immunocompetent and AM-depleted mice given luminescence below the LOQ before sacrifice and survival at 8 days above 60% in both groups. Moreover, CFU levels recovered in the lungs of AM-depleted mice treated with phage were at least four orders of magnitude lower than that in immunocompetent mice. This difference between the two groups could be detected only from CFU counts because the radiance was below the LOQ in the two groups. This difference also implies that AM directly reduce phage-induced clearance of *P. aeruginosa in vivo*. Via mathematical modeling and direct *in vivo* experiments, we suggest that macrophage-induced reduction of phage densities includes phagocytosis and other mechanisms.

Macrophages are typically classified into two functional phenotypes: pro-inflammatory M1 macrophages and anti-inflammatory M2 macrophages^39,25^. While M1 macrophages promote the inflammatory response aimed at clearance of the pathogen, including phagocytosis and recruitment of other cell types ^40,24,39,25^, the M2 macrophages participates in resolution of the infection, tissue repair and return to homeostasis ^40,24,41^. Given that at 22 h p.i the AM-depleted mice had the same bacterial load as the immunocompetent mice, we hypothesize that the observed difference in survival between these two groups of mice is related to the lack of M2 activity, which would need further experiments beyond the scope of this study. Nevertheless, prior AM depletion studies reach different conclusions on the involvement of AMs in the survival of *P. aeruginosa* infected mice ^29–31^. These differences may arise given that immune response varies depending on the bacterial strain and initial inoculum used in these studies. We identify the study by Cheung *et al.* ^30^ as the closest in terms of mouse line (BALB/c), AM depletion method and bacterial inoculum (different strain of *P. aeruginosa* however). Similar to the present findings, these authors observed no difference in CFUs in the lungs at 24 h p.i. between the AM-depleted and control groups, however the AM-depleted animals did not reach compassionate end points at 24 h p.i. The key difference is the level of AM depletion that reached 65% in their study compared to 95% in our study. This could account for the discrepancies in survival if we consider that the remaining population (35%) of AM is sufficient to recapitulate a WT phenotype.

In contrast to the contribution of AM in promoting efficient *P. aeruginosa* clearance from the lungs, we found that treatment with phage was more efficient at clearing infection in AM-depleted than in immunocompetent mice with significant lower levels of *P. aeruginosa.* Guided by a mathematical model, we discovered that phage decay *in vivo* was three times faster in immunocompetent compared to AM-depleted mice. This gap strongly points to macrophages involvement in active removal of phages from the lungs, by phagocytosis and/or other processes. There is prior evidence for phage phagocytosis by macrophages, which comes mostly from *in vitro* observations, where T2 phage was found to be internalized by rabbit macrophages and peritoneal neutrophils ^42^ and T4 phage was found inside rat dendritic cells ^43^ as well as inside murine macrophages ^44^. Interestingly, we found no significant *in vitro* phagocytosis by macrophages using the J774 macrophage cell line. However, this line originates from reticulum cell sarcoma ^45^ and not from alveoli macrophages. Here, we showed AM were able to internalize phage PAK_P1 by phagocytosis. This emphasizes that different macrophages will not necessarily respond similarly to the same stimuli. According to our mathematical model, phage decay by AM was not enough to account for the differences in CFUs and PFUs found in the lungs at end points and we suggested that an additional effect of adsorption inhibition could provide an explanation. These interactions require further investigation in order to test if AM digest phage particles and if their presence reduces adsorption rates due to spatial interference, *e.g*. “cloaking” of bacterial colonization sites. More generally, mathematical model predictions remained robust to parameter variation even insofar as a subset of assumptions and parameter values in the mathematical model could not yet be verified experimentally in this study; efforts to do so must also account for ethical considerations involved with sacrificing a much larger number of animals.

Integrating AM-induced reduction of phage densities and adsorption in a mathematical model of *in vivo* phage therapy led to bacteria, phage, and immune system population dynamics consistent with *in vivo* data from immunocompetent and AM-depleted mice. However, our study comes with caveats. We did not address phage resistance during these experiments as we did not observe a rebound in the bacterial population in phage treated groups, which is usually associated with the growth of phage resistant bacteria. However, we did incorporate resistance to phage in our mathematical model as the bacterial rebound is associated with growth of a phage-resistant bacterial population. Assumptions about phage resistance rates in the model are derived from Roach *et al*. ^14^. Hence, although we interpret the antagonistic impact of AM on phage induced lysis of *P. aeruginosa,* we recognize that evolutionary dynamics may further complicate these tripartite interactions^17^. Moreover, our model does not directly include the potential role of macrophages in murine host survival. We hypothesize that macrophage mitigate against the accumulation of damages to the tissues which can lead to the rapid decline in the health of the animal. The hypothesis that M2 macrophage are necessary for survival of *P. aeruginosa* infection requires further investigations by assessment of tissue damages for example. In the long term, indicators of tissue damages should be integrated into mathematical models to better represent the role of macrophages during infection by *P. aeruginosa* and their impact on host health and survival.

In conclusion, we have found evidence for an antagonistic interaction between phages and macrophages during phage therapy. Phages are removed faster from the lungs by AM, which potentially lower phage adsorption to bacteria as well; together these mechanisms reduce the efficacy of phage therapy in immunocompetent relative to macrophage depleted murine hosts. Since prior evidence shows that not all phages are recognized and phagocytized by the immune system, our findings suggest that selection of therapeutic phage candidates, whether as single strain or as cocktails should take into account the response of the immune system – including both neutrophils and macrophages. More broadly, identifying inclusion criteria is essential to identify subpopulations of individuals who may be responsive to given therapies – including phage therapy. The finding here, building upon prior work on the key synergistic roles of neutrophils, can help guide follow-up studies to identify actionable criteria for phage therapy use in the context of personalized medicine.

## Methods

### Animals and Ethics

BALB/c mice (nine to eleven weeks old purchased from Charles River) were housed in a pathogen-free animal facility in accordance with Institut Pasteur guidelines and European recommendations. Food and drinking water were provided *ad libitum*. Protocols were approved by the veterinary staff of the Institut Pasteur animal facility (Ref.#18.271), the National Ethics Committee (APAFIS#26874-2020081309052574 v1) and the Georgia Institute of Technology IACUC.

### Bacteria and phage

We used the bioluminescent *P. aeruginosa* strain PAKlumi that constitutively expresses the *luxCDABE* operon from *Photorhabdus luminescens* ^46^. PAKlumi was grown in Lennox Broth (LB), at 37°C with shaking at 180 rpm.

Phage PAK_P1 lysates were propagated on PAKlumi as previously described^47^. PAK-P1 was added to mid-log culture of PAKlumi at a multiplicity of infection (MOI) of 0.001. The infected culture was kept at 37°C with shaking until lysis occured. The culture was centrifuged (7000 rpm, 10 min, 4°C) and the supernatant filtered (0.2 µm) and then concentrated in Phosphate Buffered Saline (PBS) (Sigma) by ultrafiltration (Vivaflow 200™; Sartorius). The lysates were purified by cesium chloride density gradients, followed by dialysis in water and PBS. Endotoxins were removed by passages through an EndoTrap HD™ column (Hyglos, Germany) three times. Endotoxin levels were measured using the Endonext™ kit (Biomerieux) and a microplate fluorimeter (Tecan). The endotoxin level measured for the purified suspension was 10.05 EU/mL.

Bacteria from lung homogenates were enumerated by plating on cetrimide + 1 % glycerol agar plates (Sigma-Aldrich) and incubated overnight in 37°C. Phage was enumerated by spotting serial dilutions on LB agar plates that were previously inundated by a fresh culture of PAKlumi and dried under a safety cabinet 30 min before use.

### Macrophage depletion

AM depletion was obtained by the intra-tracheal administration a single dose of 400 µg of clodronate liposomes or empty liposomes (LIPOSOMA, city), four days prior infection. For the intra-tracheal administration, the mice were anesthetized by intraperitoneal injection of a mixture of two vol of ketamine (10 mg/mL) and one vol of xylazine (1 mg/mL) in a solution of NaCl 0.9%. The dose administrated to each mouse was adjusted to 0.01 mg of the ketamine-xylazine mixture per 1g of body mass.

### Flow cytometry analysis

To measure levels of AMs and PMNs three to four mice (receiving clodronate or empty lyposomes) were sacrificed and bronchoalveolar lavages (BALs) were obtained following Daubeaf et al. ^48^. Briefly, we made an incision in the trachea, inserted a 22G catheter (BD) and washed the lungs 3 times with 1 mL buffer with 2 % Fetal Bovine serum (Dulbcco’s Phosphate Buffered Saline; STEMCELL), using a 1 mL syringe (BD). Cells from broncho-alveolar lavages were filtered through a 40 mm nylon mesh and first stained with a fixable live/dead marker for 30 minutes at room temperature (RT). Cells were washed with Cell Staining Buffer (BioLegend) and stained with a cocktail of surface antibodies for 20 minutes at 4°C. Antibodies conjugated to various fluorochromes were used against the following mouse antigens: CD11b, CD11c, CD19, CD45, MHC II, Ly6C, Ly6G, F4/80, and TCRb. Cells were washed and then resuspended in 400 mL Cell Staining Buffer. Stained cells were analyzed using the ID7000 spectral cytometer (Sony). In every experiment single-stained samples in Ultracomp eBeads (Thermo Fisher), for each individual fluorochrome, were used as control. Data were analyzed using FlowJo software (BD).

### Estimating depletion effect of clodronate

We used the following model to represent clodronate pharmacokinetics and its effect on AM:

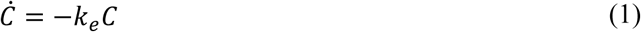

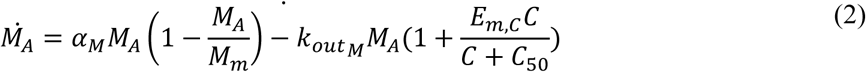

Where *C* (Eq. 1) is the clodronate amount in the lung, *k*_*e*_ is the clodronate elimination rate from the lung, *M*_*A*_ (Eq. 2 is the AM concentration, α_*M*_ the self-renewal rate of macrophages, *M*_*m*_ the maximal concentration of macrophages, *k*_*outM*_ the elimination rate of macrophages, *E*_*m*,*C*_ the maximal effect of clodronate and *C*_50_ the clodronate amount to reach 50% of the maximal effect. The initial condition of *C* is given by the intratracheal dose of clodronate. The initial condition of *M*_*A*_ is given by the data from uninfected, untreated and non-depleted mice, assuming the empty liposomes did not affect the alveolar macrophage level 96 h after administration.

### Murine model of acute pneumonia

Mice were first anesthetized by intraperitoneal injection of the same mixture of ketamine and xylazine as described above at a dose of 0.008 mg per 1 g of body mass. An exponentially growing culture of PAKlumi was washed three times in PBS before its density was adjusted to 1.25 x 10^8^/mL in PBS. Mice were infected with a single intranasal administration of 40 µL (5 x 10^6^ CFU). Phage treatment was administered intranasally at a dose of 5 x 10^7^ plaque forming units (PFU; MOI 10) in 40 µL PBS at 2 h p.i.. The progression of infection was monitored by measuring the luminescence (IVIS, Perkin Elmer) produced by the bacteria from the lungs area of mice that were anesthetized by isoflurane inhalation. The LivingImage software (version 4.7.3; Xenogen) ^14,34^ was used to collect emitted photons within a consistent square ROI (2.5 by 3.0 cm), corresponding to the size and localization of the lungs. At end points lungs were collected and kept on ice in C tubes (gentleMACs^TM^, Miltenyi Biotec) containing 1 mL PBS before homogenization with a tissue dissociator (Oligo-Macs, Miltenyi Biotec). The samples were serially diluted and plated on cetrimide agar.

### Conversion of bioluminescence data into bacterial load in lungs

We established the correlation between bioluminescence and CFU count using data from PAKlumi-infected and untreated animals for which emitted photons were recorded before lungs were taken, homogenized and plated to count bacteria (Sup. Fig. 3). The relationship obtained allowed us to derive from the luminescence signal an equivalent counts of CFU per mL of lung (noted CFUeq/mL) that was used to obtain the bacterial loads in all infected animals (untreated or treated by phages) over time.

### Estimating *in vivo* AM-associated phage decay

Two groups of uninfected mice (AM-depleted and EL control) received a single dose of 5 x 10^7^ PFU of phage PAK_P1 administered intranasally. At 24, 48, 72, and 96h after phage administration, mice were sacrificed lungs were recovered, homogenized, and phages counts were measured as described above. The phage decay was described using a monoexponential decay in AM-depleted mice (Eq. 3):

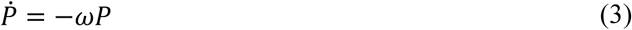

where ω is the decay rate, which was estimated from uninfected and AM-depleted mice to obtain the decay of the phage PAK_P1 in absence of macrophages (assuming that the residual amount of macrophages was negligeable). As the initial dose of phage (*d*_*P*_, in PFU) was known and the initial phage concentration was given by the fit, the scaling factor (or distribution volume) *f*_*P*_ was estimated, with the initial condition given as *P*(0) = *d*_*P*_/*f*_*P*_. The phage decay rate was augmented as follows by interactions with AM (Eq. 4):

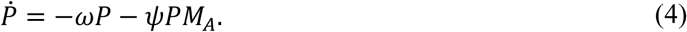

The AM-associated decay rate parameter ϕ was estimated from uninfected and control (empty liposome) mice, with ω fixed from experiments with AM-depleted mice. The AM level (*M*_*A*_) was considered as stable during all experiments in the control group. Estimations were performed by naïve pooling of data (*i.e.*, considering all the measures for each group (with or without clodronate) as if they came from a single individual and performing a non-linear regression, Fig. 2B).

### Quantification of phage DNA from AM-sorted from bronco-alveolar lavages

Anesthetized mice (see above) were intranasally administered with either a single dose of 5 x 10^7^ PFU of phage PAK_P1 or PBS (40 µL). Four hours later, mice were euthanized, and bronchoalveolar lavage (BAL) was conducted according to Busch et al.^49^. Lavage was performed using 10 times 1 mL (total volume = 10 mL) of D-PBS containing 2% FBS (STEMCELL) and 2 mM EDTA (Invitrogen), which was pre-heated to 37°C. A volume of 40 µL of BALs was taken to measure the number of viable phages while the rest was centrifuged at 500 g for 10 minutes at 4°C, and the supernatant was discarded to remove free phage particles. The cells were then resuspended in hemolysis buffer (Biolegend) for 5 minutes at room temperature to lyse erythrocytes, followed by another centrifugation step (which correspond to a washing step to remove further any remaining free phage particles). Subsequently, cells were stained with antibodies against CD45 (BD), CD11c (Miltenyi Biotec), and Singlec-F (BD) (diluted 1:200) for 20 minutes in the dark at room temperature. After addition of 500 µL of D-PBS, AM were identified as positive for all three markers and sorted using the FACSAria III cell sorter (BD) in to 1 mL. Then 2 µL of each sample was used as template for a qPCR reaction with 450 nM of each primer (F-ACGCCAGACCGAAGACAACT and R-CGTTCCAATCTCCCGCAACG) targeting the terminase large subunit (Gene ID: 10351274) using the PowerSYBR Green PCR master mix (Applied Biosystems) in a final volume of 15 µL and run on BioRAD CFX Opus 96 Real-Time PCR System (denaturation step at 95°C for 20 minutes followed by 40 cycles of amplification with denaturation at 95°C for 10 seconds and annealing/elongation at 55°C for 60 seconds). Fluorescence (FAM; Ex/Em 465/510nm) was measured at the end of each cycle, and melting curve analysis was conducted to verify the specificity of the amplified PCR product. The number of DNA copies in each reaction was determined by comparing the Cycle threshold (Ct) to that of a sample with a known DNA concentration using a standard curve. Genomic DNA for standard curves was extracted using the Wizard Plus DNA purification system (Promega). DNA concentrations in ng/mL were measured by absorbance at 260 nm using a spectrometer (Eppendorf) and converted to gene copies/mL by inputting the genome length and concentration into the Technology Network Copy Number Calculator (https://www.technologynetworks.com/tn/tools/copynumbercalculator) for precise determination of copy numbers. The number of gene copies/mL was divided by the number of AM/mL sorted.

### Phage decay assessment in presence of J774 macrophages

Macrophage cell line J774.A1 (American Type Culture Collection No. TIB67) derived from BALB/c mice was maintained in RPMI 1640 medium + Glutamax™ (Gibco, Life Technologies Corporation, Grand Island, USA) and supplemented with 10 % (v/v) fetal calf serum (Eurobio scientific, Les Ulis, France). Cells were seeded at a density of 2 x 10^5^ cells per well in 24-well tissue culture plates and grown for 18 h at 37°C in an atmosphere enriched with 5% CO_2_. Phage PAK_P1 was added to J774 monolayers or RPMI medium (control wells) at different phage:cells ratios ranging from 1:1 to 1000:1. At several times points (0, 1h, 2h, 4h, 6h and 24h) supernatants were collected and phage titration performed as described above.

### Phagocytosis assay of phage PAK_P1 in presence of AM

BALs from 10 uninfected mice were collected according to Busch et al.^49^. Lavages were performed using 5 times 1 mL (total volume = 5 mL) of D-PBS containing 2% FBS (STEMCELL) and 2 mM EDTA (Invitrogen), which was pre-heated to 37°C. The cells were then filtered through a 70 µm cell strainer and collected by centrifugation (400g, 5 min, room temperature). Next, 1 ml of red blood cell lysis buffer (BioLegend) was added, followed by a 5-minute incubation at room temperature. The lysis was stopped by adding 5 ml of complete medium (RPMI 1640, 1X GlutaMax, 10% FBS, 1X PenStrep, 1X sodium pyruvate) and cells were collected following a centrifugation (400g, 5 min, room temperature) and resuspended in a total volume of 10 mL of complete medium. AMs (300 µL) were then distributed in chambers (Lab-Tek II Chamber Slide 8-wells, ThermoFisher) and incubated with or without 20 µM Cytochalasin B (C2743-200UL, Sigma-Aldrich) for 30 minutes at 37°C. Phage PAK_P1 or beads (0.2 µm FluoSpheres™ Sulfate-Modified Yellow-Green Microspheres, Invitrogen #F8848) were added at a ratio of approximately 1000 phages or beads per AM. After 1, 3 and 5 h of incubation at 37°C 4% PFA was added and “slides” were kept overnight at 37°C. To detect phage particles a guinea pig primary antibody raised against phage PAK_P1 (Covalab) was applied (1/100 dilution), followed by a secondary antibody (anti-guinea pig Alexa Fluor 488, Abcam 150185, 1/500 dilution). The cytoplasm was stained with the CellMask membrane stain (C10046 ThermoFisher, 1/10,000), actin was labeled with Phalloidin 546 (A-30106 ThermoFisher, 1/750), and the nucleus was stained with DAPI (D1306 ThermoFisher, 1/8,000). Images were acquired on IX81 inverted microscope. A set of n=13 to 47 macrophages per condition were randomly selected and analyzed (see Sup. Methods and Sup. Fig. 6).

## Model building, parameter estimation and model evaluation

The final mathematical model described in detail in Sup. Methods was built following a sequential approach using the estimated parameters of clodronate effect on macrophages from the uninfected untreated mice and the macrophage effect on phage decay from the uninfected phage-treated mice. Key parameters of phage-immune cells-bacteria interactions were estimated from all the infected mice (AM-depleted and control, phage-treated and untreated). Inference procedures were performed in a non-linear mixed effect model framework using the SAEM algorithm implemented in Monolix software version 2021R2 ^50^. The individual parameters θ_*i*_ (with i the individual) followed a log Normal distribution (θ_*i*_ = μ exp(η_*i*_), with μ the median parameter and η_*i*_ the random effect for *i*, η_*i*_ following a Normal distribution of expectancy 0 and standard deviation ω_*i*_). All inference procedures assumed an additive Gaussian error model on the log-transformed (bacteria and phage) data and a proportional error model on neutrophils (lung and blood), monocytes and macrophage data. The details of the choice and precision of estimation of parameter values (data used for estimation or reference in case of fixed parameter from the literature) are given in Table S1. The model selection was based on Bayesian information criterion ^51^. Individual fits and residuals were investigated to evaluate the different models. The final model was also evaluated using normalized prediction distribution errors ^36^. Graphic representations and simulations from the model and estimates were made with R using the RsSimulx package (version 2.0.2).

### Statistical analysis

Statistical analyses were performed using the Prism software (GraphPad, v9.5.0). Group comparisons were done using the Mann Whitney U test (independent samples), or the Wilcoxon sign test (paired samples). Log-rank (Mantel-Cox) test was used to compare survival curves. Pearson r correlation and simple line regression were used for correlation and dependency assessments.

## RESOURCE AVAILABILITY

### Lead contact

Further information and requests for resources and reagents should be directed to and will be fulfilled by the lead contact, Laurent Debarbieux.

### Materials availability

The bacterial strain PAKlumi and phage PAK_P1 are available from the lead contact upon completion of a Materials Transfer Agreement (MTA).

### Data and code availability

The source code for the mathematical model is included in the supplemental files as Data S1 and is available in GitFront: https://gitfront.io/r/jeremyseurat/c5QMQsU2Kjj3/MacrophagePhage/

The raw data is available at: https://entrepot.recherche.data.gouv.fr/privateurl.xhtml?token=1fadeb85-b662-4b62-9ff3-ebeef9cc64ab

## Acknowledgments

We thank Tanguy Dequidt and Baptiste Gaborieau for sharing data used for the calibration curve. We thank Sandrine Schmutz from the Cytometry platform of Institut Pasteur for performing the flow cytometry analysis. We thank Elisa Gomez Perdiguero (Institut Pasteur), Thomas Secher (CEPR INSERM U1100) and Molly Ingersoll (Institut Pasteur, Institut Cochin) for helpful discussions during revisions. We thank Stéphane Rigaud and Jean-Yves Tinevez from the Image Analysis Hub of Institut Pasteur. This study was supported by grants from the National Institutes of Health (1R01AI46592-01 to JSW and LD), from the Conseil Régional Ile de France Chaire Blaise Pascal (to JSW) and from Agence Nationale de la Recherche (ANR-19-AMRB-0002 to LD).

## DECLARATION OF INTERESTS

The authors declare no competing interests.

## AUTHOR CONTRIBUTIONS

Conceptualization and methodology, J.S., J.S.W., L.D., Q.B.-D. and S.Z.; investigations, C.M., C.N.M.N., E.E., J.M., J.S., M.T., Q.B.-D., R.R.-G., S.E. and S.Z.; formal analysis, J.S. and S.Z.; supervision and funding acquisition, J.S.W. and L.D.; writing—original draft, J.S. and S.Z.; writing—review and editing, J.S., J.S.W., L.D. and S.Z.

## Supplementary material

**Supplementary Figure 1.**
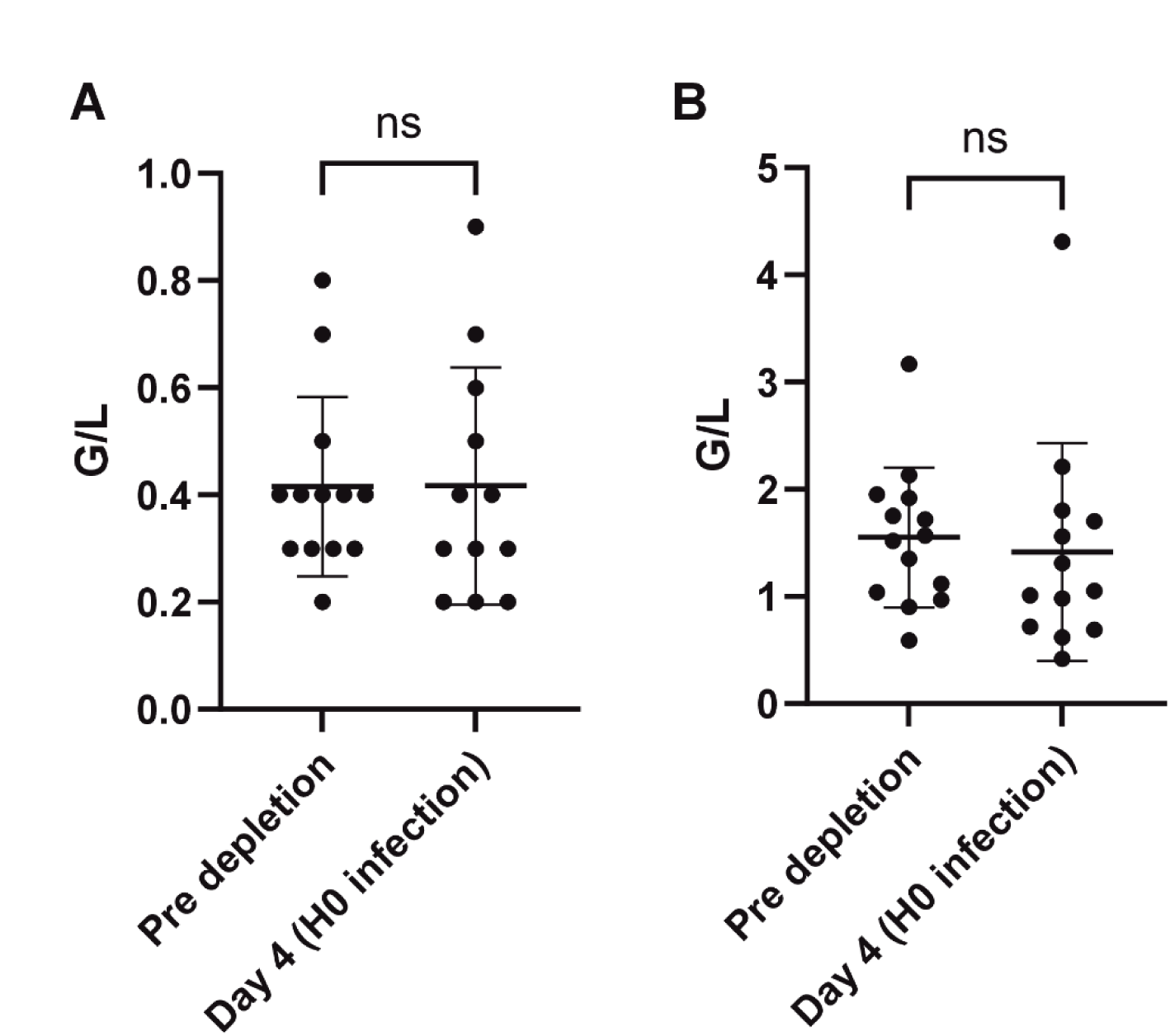
Levels of immune cells in blood from AM-depleted mice. Blood levels of (A) monocytes (B) neutrophils in mice treated before (n=13 monocytes) and four days post clodronate administration (n=12) to deplete AM (Wilxocon signed rank test was used for group comparisons).

**Supplementary Figure 2.**
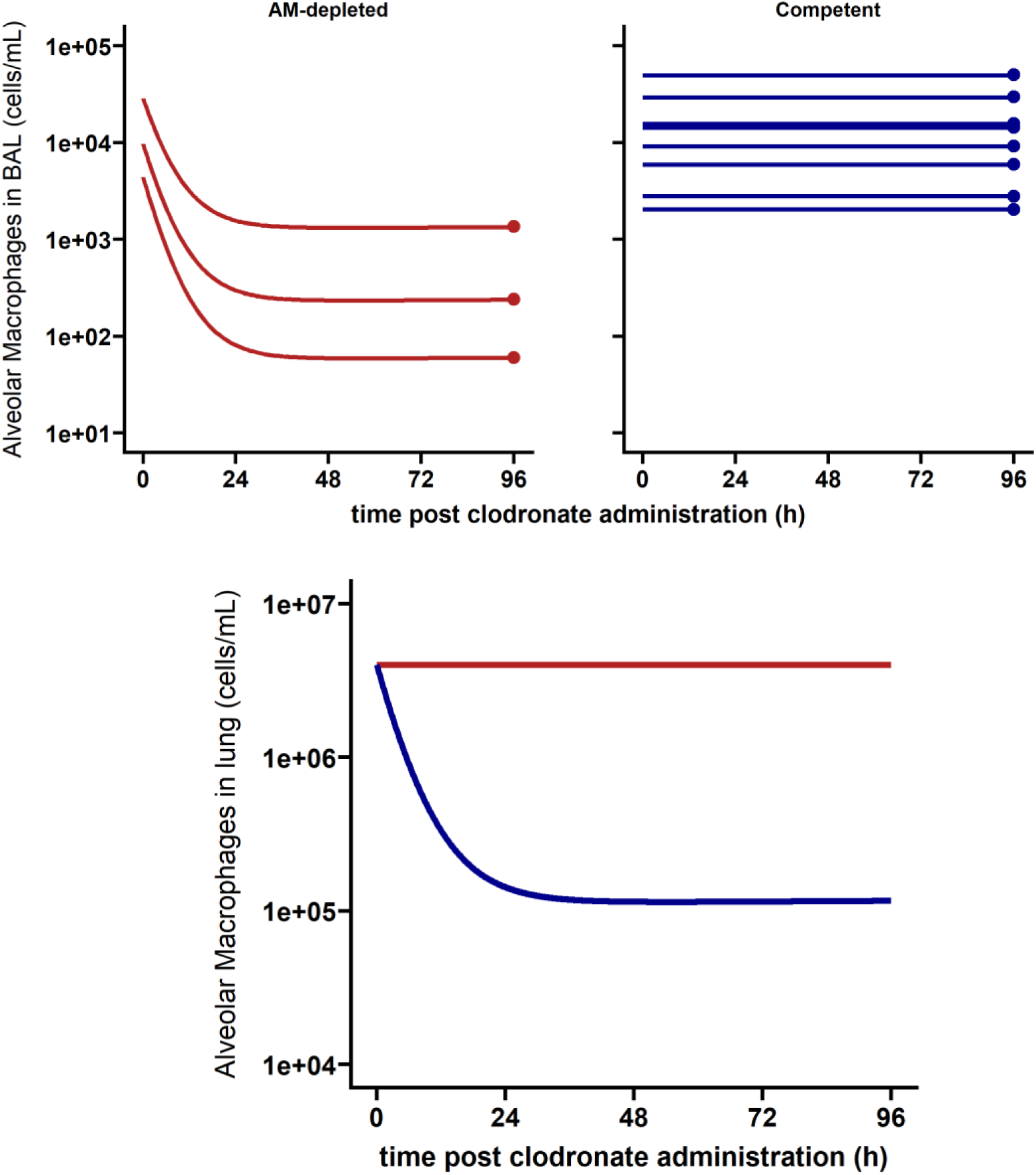
Clodronate effect on alveolar macrophages (AM). The individual data in the broncho-alveolar lavage (dots) and individual model fits (see Eq. 1-2 and parameters in Sup. Methods) are represented in uninfected and immunocompetent (empty liposome) mice in red (top left, n=8) and in uninfected and AM-depleted mice (top right, n=3). The model-based median dynamics of AM in the lung is shown at the bottom for these two groups.

**Supplementary Figure 3.**
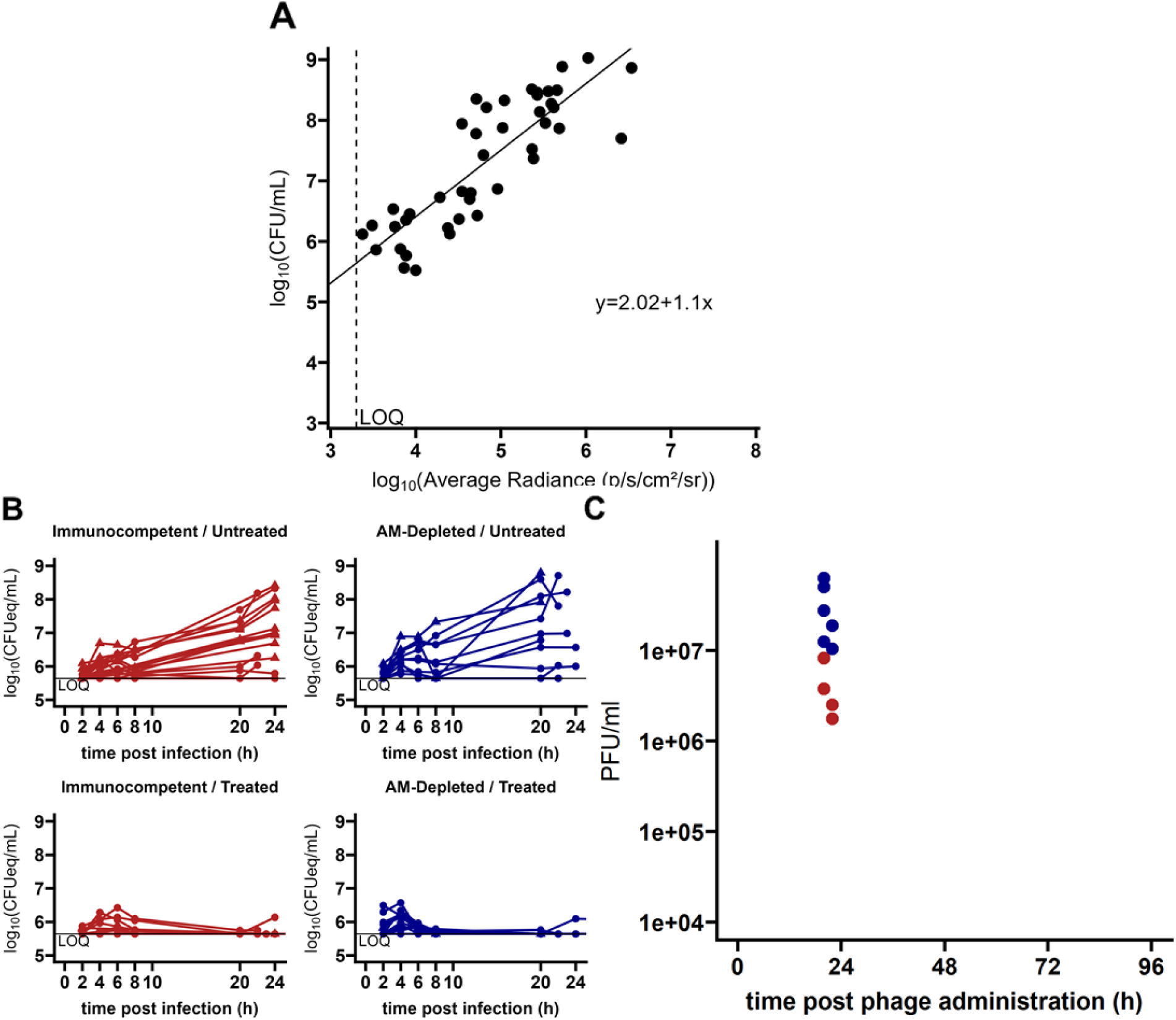
Correlation of average radiance and CFU, individual *P. aeruginosa* and phage loads in mice lungs. (A) The regression line equation is *log*_10_(*CFU*/*mL*) = 2.02 + 1.10 *log*_10_(*p*/*s*/*cm*^2^/*sr*). The adjusted coefficient of determination (adjusted-R²) is 0.71 (p<10 ^-3^, Pearson’s correlation test). The dashed vertical line represents the lower limit of quantification (LOQ) of 2000 p/s/cm²/sr. Data were collected from BALB/c mice (n=44) infected by the same dose of strain PAKlumi, from experiments previously carried out in our laboratory independently of this study. (B) The individual bacterial load (converted in CFUequivalents/mL from the correlation shown in panel A are represented in the four subpanels from AM-depleted and immunocompetent groups of infected mice untreated or phage-treated. The dots from each individual are connected by a line. The horizontal black line represents the lower limit of quantification (LOQ). (C) Phage density in the lungs at sacrifice 20 to 22h after phage administration (phage were administered 2 hours after infection) in immunocompetent (EL, red dots) and AM-depleted (blue dots).

**Supplementary Figure 4.**
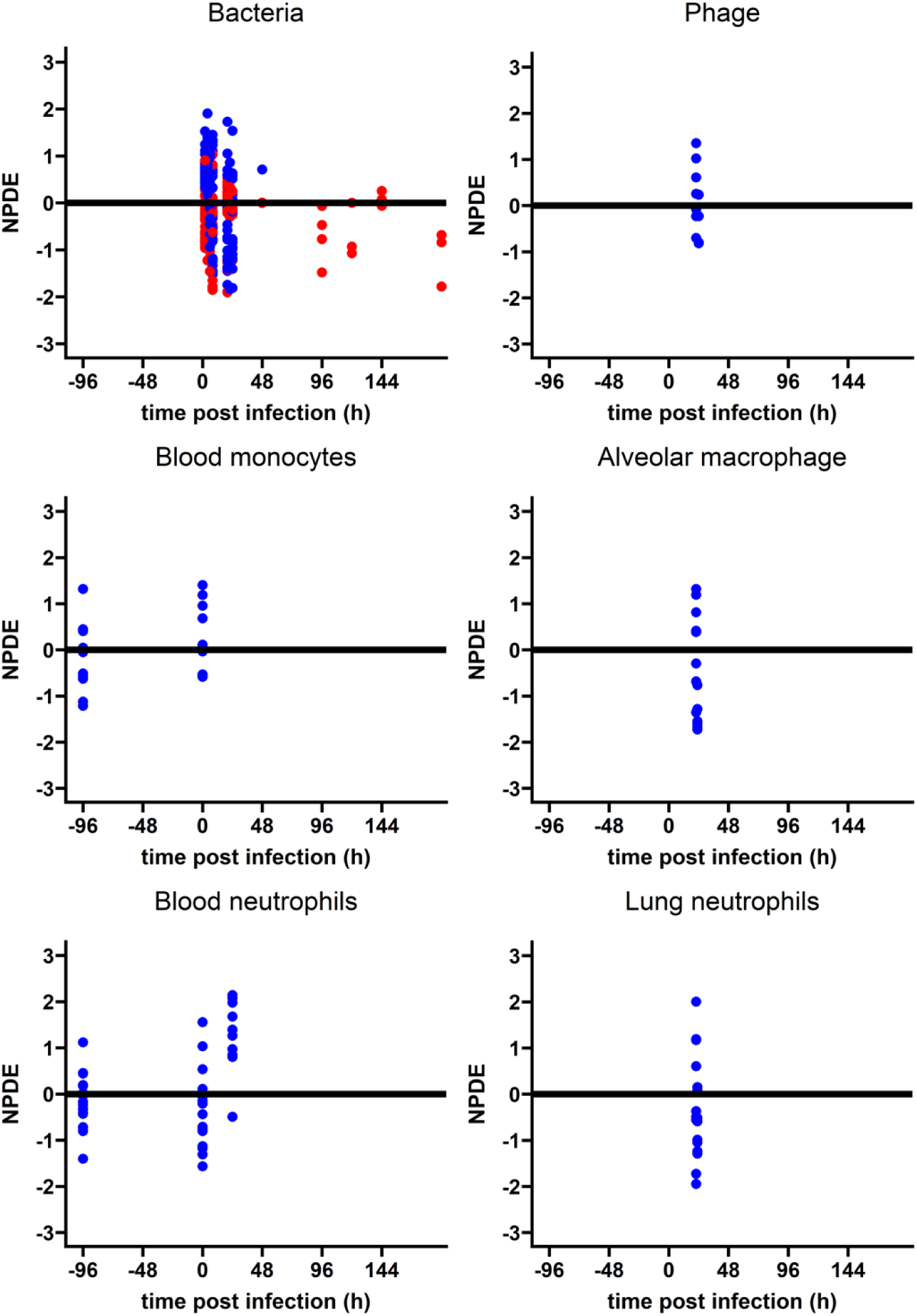
Model evaluation: normalized predictions distribution errors (NPDE). Each dot was included in the data set used in the inference from the final model (Sup. Methods). Blue dots represent quantified observations and red dots represent below the limit of quantification observations.

**Supplementary Figure 5.**
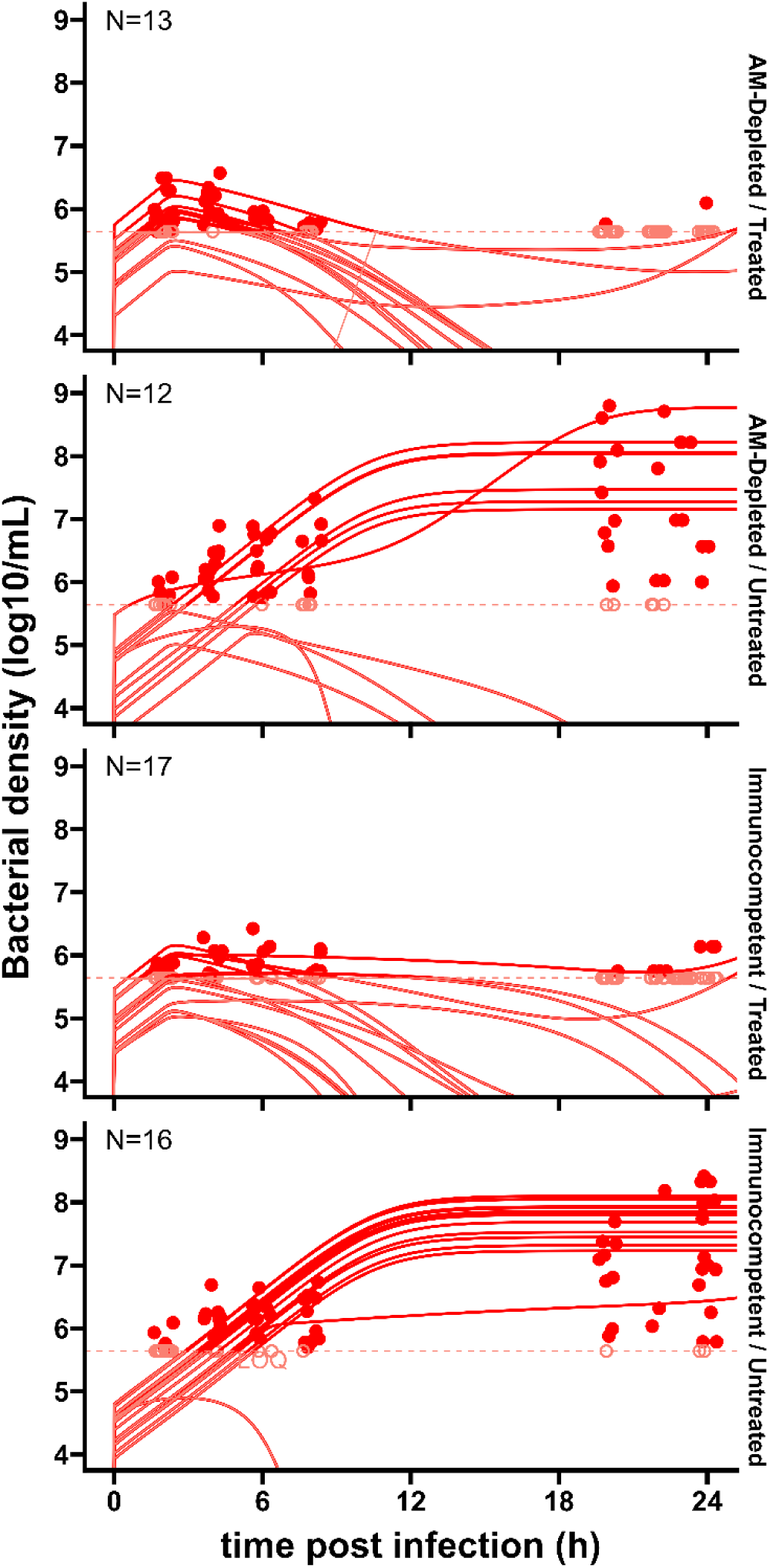
Longitudinal bacterial density and model fits from all infected mice (N=58). Lines represent model-based individual predictions of bacterial density in the four mice groups separated vertically into different panels, with the number of mice in each group indicated at the top left of each panel. Dots are observed bacterial densities. The light red horizontal dashed lines represent the limit of quantification (LOQ) for bacterial density (LOQ = 5.6 log10 CFUeq/mL), light red circles crossed by this line at their center are below the LOQ of bacterial density and light red lines are below the LOQ of bacterial load model-based predictions (i.e. the discrepancy between data and model cannot be evaluated from this plot in case of below LOQ data).

**Supplementary Figure 6.**
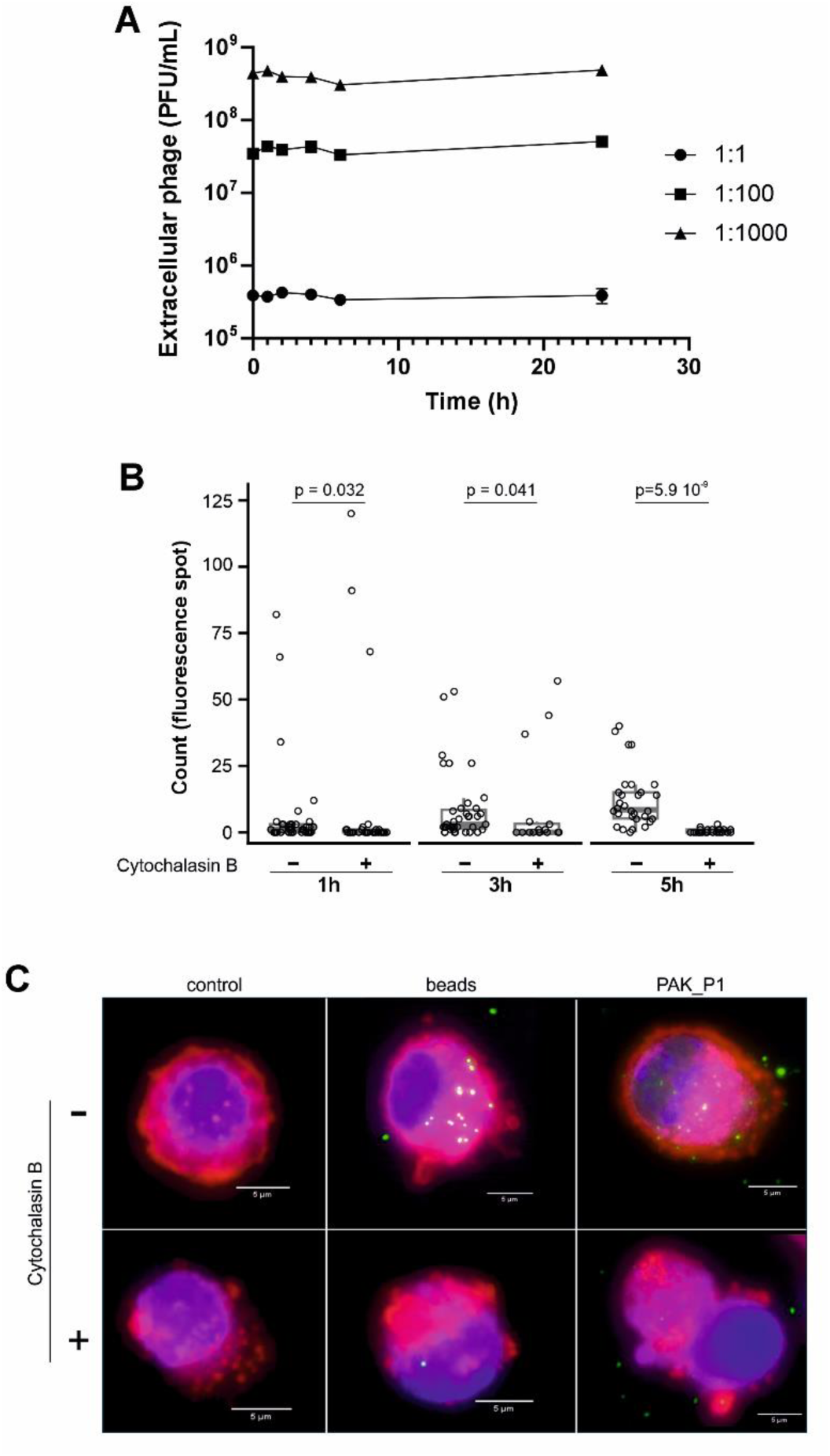
***In vitro* interactions between macrophages and phage PAK_P1. A.** J774 BALB/c derived macrophages were mixed with PAK_P1 at a ratio of macrophage: phage of 1:1, 1:100 and 1:1000. Extracellular phage was sampled out at the indicated time intervals and quantified by spot assay. Group comparison (Mann-Whitney U test) between 0, 6 and 24 h showed no significant differences. n=3. **B.** Counts of fluorescent signals from beads in individual AM collected from uninfected mice and incubated without (control) or with beads in absence or presence of cytochalasin B at indicated time points (n=16 to 40 for each condition). **C.** Representative images of individual murine AM with fluorescent signals associated to either control, beads or phage PAK_P1 after 3 h of incubation (AM:phage and AM:beads ratios of 1:1000).

**Supplementary Figure 7.**
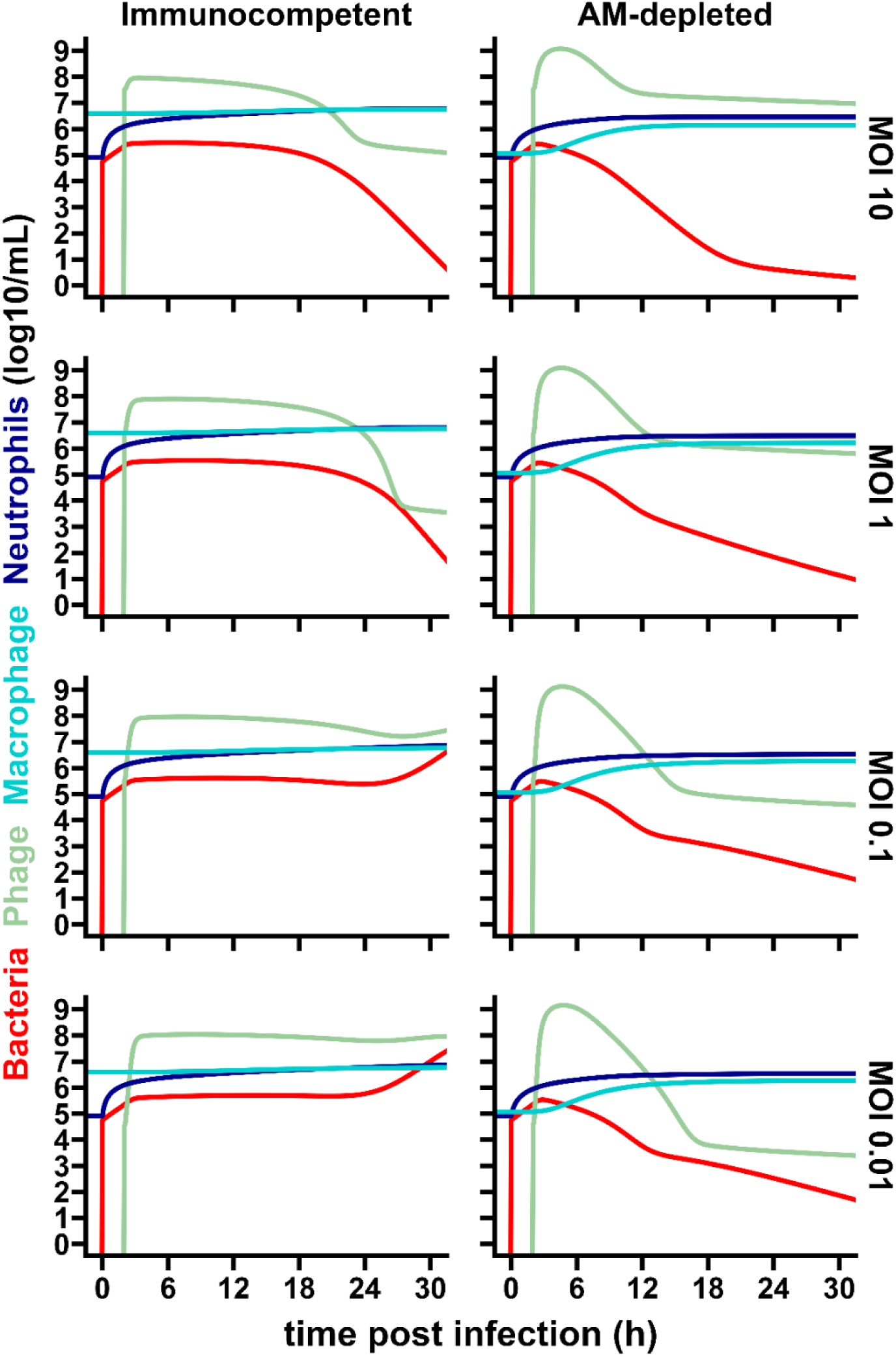
Median dynamics cell and phage populations following different phage doses. Based on the model (Fig. 2, Eq S1-S15) and its median parameters (Sup. Methods) median dynamics of bacteria, macrophages, neutrophils and phage were estimated for four treatment conditions ranging from multiplicity of infection (MOI) 10 (*i.e.*, the conditions of the experiments) to MOI 0.01.

## Notes

### Competing Interest Statement

The authors have declared no competing interest.

### Summary of Updates

Addition of an experiment showing the phagocytosis of phages by alveolar macrophages.

