## Supplementary Methods for "Macrophage-induced reduction of bacteriophage density limits the efficacy of *in vivo* pulmonary phage therapy"

### Deconvolution, image processing and analysis of phagocytosis assays with alveolar macrophage

All images were deconvolved with Huygens Professional version 24.04 (Scientific Volume Imaging, The Netherlands, `http://svi.nl`), using the CMLE algorithm, with Acuity: 26.3, SNR: 47.5, max 100 iterations, and background automatically estimated with an estimation radius of 0.7 µm. Deconvoluted images where first rescaled along the z-axis to correct for the pixel anisotropy and cropped of 1.5 um from the top and from the bottom to remove possible deconvolution artefacts. Cell contour mask where delimited by hand using Fiji in 2D and propagated in 3D to fit the image shape. The phages and beads where then detected using a Difference of Gaussian ($\sigma_1 = 2$, $\sigma_2 = 4$) followed by a local maxima detector filter. The detected peaks are then filtered based on their quality (normalized DoG intensity $> 0.75$), inside the cell contour mask, and having an GFP intensity superior to a predetermined threshold (t=22000) computed based on the maxima background intensity of the eGFP in a control experiment. The processing script is done in python (3.12) and uses the following scientific libraries: scikit-image (0.24.0) [S13], pyclesperanto (0.16.1) [S9], scipy (1.14.1) [S14], and pandas (2.2.3) [S8].

**Mathematical model**

A mathematical model was built to recapitulate the interactions between bacteria, phage, neutrophils and alveolar macrophages (AM) in the lung during infection. The "immunophage synergy" model as presented in Roach et al. [S10] showed the synergy between neutrophil and phages to kill bacteria. In the model presented here, the role of neutrophils and phages is still included, in addition to this, we show the interactions of macrophages with bacteria, phages and neutrophils (Fig. 2, Eq. S1-S15 below).

*Clodronate*

$$\dot{A_C} = \overbrace{-k_{e_C} A_C}^{\text{clodronate lung clearance}}, \tag{S1}$$

*Blood neutrophils*

$$\dot{N_B} = \overbrace{R_{in_N}}^{\text{production}} \overbrace{-k_{out_N} N_B}^{\text{elimination}} \overbrace{-r_{max} N_B \frac{B}{B + K_{N_n,B}} \left(1 - I_{k_M}\left(1 - \frac{M_A}{M_A + K_{N_n,M}}\right)\right)}^{\text{neutrophil recruitment}}, \tag{S2}$$

*Lung neutrophils*

$$\dot{N_L} = \overbrace{r_{max} N_B \frac{B}{B + K_{N_n,B}} \left(1 - I_{k_M}\left(1 - \frac{M_A}{M_A + K_{N_n,M}}\right)\right) \frac{V_b}{V_l}}^{\text{neutrophil recruitment}}, \tag{S3}$$

*Blood monocytes*

$$\dot{M_B} = \overbrace{R_{in_M}}^{\text{production}} \overbrace{-k_{out_M} M_B}^{\text{elimination}} \overbrace{-k_{M_{bl}} M_B \left(1 + \frac{r_{max} - k_{M_{bl}}}{k_{M_{bl}}} \frac{B}{B + K_{N_m,B}}\right) + k_{M_{lb}} M_L \frac{V_l}{V_b}}^{\text{monocyte recruitment and exchanges with lung}}, \tag{S4}$$

*Lung monocytes*

$$\dot{M_L} = \overbrace{k_{M_{bl}} M_B \left(1 + \frac{r_{max} - k_{M_{bl}}}{k_{M_{bl}}} \frac{B}{B + K_{N_m,B}}\right) \frac{V_b}{V_l} - k_{M_{lb}} M_L}^{\text{monocyte recruitment and exchanges with lung}} \overbrace{-d_M \left(B/\left(B + K_{N_m}\right)\right) \left(1 - \frac{M_A}{M_{max}}\right) M_L}^{\text{macrophage differentiation}}, \tag{S5}$$

*Immature macrophages*

$$\dot{M_{LT_1}} = \overbrace{d_M (B/(B + K_{N_m})) \left(1 - \frac{M_A}{M_{max}}\right) (M_L - M_{LT_1})}^{\text{macrophage differentiation, stage 1}}, \tag{S6}$$

$$\ldots \tag{S7}$$

$$\dot{M_{LT_5}} = \overbrace{d_M (B/(B + K_{N_m})) \left(1 - \frac{M_A}{M_{max}}\right) (M_{LT_4} - M_{LT_5})}^{\text{macrophage differentiation, stage 5}}, \tag{S8}$$

*Alveolar macrophages*

$$\dot{M_A} = \overbrace{\alpha_{AM} M_A \left(1 - \frac{M_A}{M_{max}}\right)}^{\text{macrophage self-renewal}} \overbrace{-k_{out_{AM}} M_A \left(1 + \frac{E_{max,C} A_C}{A_C + A_{50,C}}\right)}^{\text{decay and clodronate effect}} \overbrace{+d_M (B/(B + K_{N_m})) \left(1 - \frac{M_A}{AM_{max}}\right) M_{LT_5}}^{\text{macrophage differentiation}}, \tag{S9}$$

*Susceptible bacteria*

$$\dot{S} = \overbrace{\alpha_S S \left(1 - \frac{B}{B_{max}}\right)}^{\text{growth}} \overbrace{-\frac{\varepsilon_N N_L S}{1 + B/K_{D,N}}}^{\text{neutrophil killing}} \overbrace{-\frac{\phi(1 - \iota_M M_A^{\gamma_M}/(M_A^{\gamma_M} + M_{50}^{\gamma_M}))S(P + P_n)}{1 + (P + P_n)/P_C}}^{\text{phage adsorption}}, \tag{S10}$$

*Resistant bacteria*

$$\dot{R} = \overbrace{\alpha_R R\left(1-\frac{B}{B_{max}}\right)}^{\text{growth}} \overbrace{-\frac{\varepsilon_N N_L R}{1+B/K_{D,N}}}^{\text{neutrophil killing}}, \tag{S11}$$

*Infected and latent bacteria*

$$\dot{I}_1 = \overbrace{\frac{\phi\left(1-\iota_M M_A^{\gamma_M}/\left(M_A^{\gamma_M}+M_{50}^{\gamma_M}\right)\right) S(P+P_n)}{1+(P+P_n)/P_C}}^{\text{phage adsorption}} \overbrace{-\frac{\varepsilon_N N_L I_1}{1+B/K_{D,N}}}^{\text{neutrophil killing}} \overbrace{-eI_1}^{\text{latency}}, \tag{S12}$$

*Infected and productive bacteria*

$$\dot{I}_2 = \overbrace{eI_1}^{\text{latency}} \overbrace{-\frac{\varepsilon_N N_L I_2}{1+B/K_{D,N}}}^{\text{neutrophil killing}} \overbrace{-eI_2}^{\text{lysis}}, \tag{S13}$$

*Administered bacteriophage*

$$\dot{P} = -\overbrace{\omega P}^{\text{decay}} \overbrace{-\frac{\phi(1-\iota_M M_A^{\gamma_M}/(M_A^{\gamma_M}+M_{50}^{\gamma_M}))(S+I_1+I_2)P}{1+(P+P_n)/P_C}}^{\text{adsorption}} \overbrace{-M_A\psi P}^{\text{AM-mediated decay}}, \tag{S14}$$

*Bacteriophage from bacterial lysis*

$$\dot{P}_n = \overbrace{\beta e I_2}^{\text{burst}} - \overbrace{\omega P_n}^{\text{decay}} \overbrace{-\frac{\phi(1-\iota_M M_A^{\gamma_M}/(M_A^{\gamma_M}+M_{50}^{\gamma_M}))(S+I_1+I_2)P_n}{1+(P+P_n)/P_C}}^{\text{adsorption}} \overbrace{-M_A\psi_n P_n}^{\text{AM-mediated decay}}. \tag{S15}$$

This system of nonlinear ordinary differential equations includes the following variables: $A_C$ the clodronate quantity (mg) in the lung (Eq. S1), $N_B$ the neutrophil concentration (cells/mL) in the blood (Eq. S2), $N_L$ the neutrophil concentration (cells/mL) in the lung (Eq. S3), $M_B$ the monocyte concentration (cells/mL) in the blood (Eq. S4), $M_L$ the monocyte concentration (cells/mL) in the lung (Eq. S5), $M_{LT_n}$ the monocyte to macrophage transit compartments (cells/ml) in the lung (Eq. S6-S8), $M_A$ the alveolar macrophage concentration (cells/mL) in the lung (Eq. S9), $S$ the susceptible bacteriaquantity (CFUeq) in the lung (Eq. S10), $R$ the resistant bacteria quantity (CFUeq) in the lung (Eq. S11), $I_1$ the infected and latent bacteria quantity (CFUeq) in the lung (Eq. S12), $I_2$ the infected and producer bacteria quantity (CFUeq) in the lung (Eq. S13), $P$ the administered phage quantity (PFU) in the lung (Eq. S14), $P_n$ the phage quantity (PFU) from bacterial lysis (Eq. S15).

In this model, the clodronate ($A_C$) decays linearly and depletes alveolar macrophages ($M_A$. Depending on the bacterial density, lung neutrophils ($N_L$) are recruited from the blood ($N_B$) at a maximal recruitment rate $r_m$. In the same way, lung monocytes ($M_L$) are recruited from the blood ($M_B$). Monocytes differentiate into alveolar macrophages within lung tissue via 5 compartments ($M_{LT_1}$, ..., $M_{LT_5}$) allowing a delay between infection and the onset of macrophage increase, as observed in the study by Koogushi et al. [S3]. The importance of macrophage involvement in neutrophil recruitment during infection is captured by the parameter $I_{k_M}$. Bacteria are divided into different populations: susceptible ($S$) and resistant ($R$) to phages. Both are able to grow (at a rate $\alpha_S$ or $\alpha_R$, respectively) and to be killed by $N_L$ (at a maximal rate of $\varepsilon_N$ and with $K_D$ the bacterial quantity at which the half of $\varepsilon_N$ is reached to account for immune evasion). $S$ can be infected by phages ($P$ and $P_n$, respectively) following a maximal adsorption rate $\phi$ to become infected and latent ($I_1$), then infected and producer ($I_2$) which are lysed to release $\beta$ new phages ($P_n$). This model also accounts for superinfection (i.e. phages can infect $S$, $I_1$ and $I_2$). In this model, the resistant population is completely resistant to phage (i.e. $R$ cells cannot enter into the lytic cycle). The total number of bacteria ($B$) is the sum of $S$, $R$, $I_1$ and $I_2$ ($B = S + I_1 + I_2 + R$). $P$ and $P_n$ are cleared by macrophages at respective different rates $\psi$ and $\psi_n$ to take into account the theory that macrophages are more present where bacteria, bacterial debris, damaged tissue and new phages from lysed bacteria are located than in other lung parts. As in Roach et al. [S10], the model accounts for an adsorption saturation term $P_C$, which constrains the adsorption rate at the maximal level $\phi$ if phage are in high density. To account for observed differences in both bacterial and phage densities in competent vs. AM-depleted mice, the adsorption is also reduced at a maximal percentage of $i_M$ depending on the macrophage level, the sigmoidicity of this effect being governed by the $\gamma_M$ parameter. A summary of all the model parameters units, values and signification is given in Table S1.

TABLE S1: Estimated parameters of the phage-bacteria-macrophage-neutrophils model

| Parameter | Unit | Value (RSE) [$\omega$] | Meaning | From |
|---|---|---|---|---|
| $k_{e_C}$ | /h | 0.14 | Linear elimination rate of clodronate | Thepen et al. [S12] |
| $N_{B_0}$ | cells/mL | 1.8 $10^6$(12)[0.41] | Blood neutrophils at baseline | This study (uninfected) |
| $k_{out_N}$ | /h | 0.115 | Elimination rate of blood neutrophils | Boxio et al. [S1] |
| $R_{in_N}$ | cells/mL/h | 2.0 $10^5$ | Production rate of blood neutrophils | $N_{B_0} \times k_{out_N}$ |
| $V_b$ | mL | 1.2 | Blood volume of a mouse | Mitruka et al. [S5] |
| $N_{L0}$ | cells/mL | 81 500 [0.66] | Lung neutrophils at baseline | This study (uninfected) |
| $f_{cN}$ | mL(lung)/mL(BAL) | 0.004 | Correction factor for lung neutrophils | This study (uninfected) |
| $V_l$ | mL | 1 | Lungs volume of a mouse | Irvin et al. [S2] |
| $r_{max}$ | /h | 0.48 [0.6] | Neutrophil recruitment rate | This study* |
| $K_{N_n,B}$ | CFUeq | 1.4 $10^6$ | Bacteria for half of max recruitment | Roach et al. [S10] |
| $I_{k_M}$ | cells/mL | 0.5 | Fraction of recruitment due to macrophage | Koogushi et al. [S3] |
| $K_{N_n,M}$ | cells/mL | 1 $10^6$ | Macrophages for half of max recruitment | This study* |
| $M_{B_0}$ | cells/mL | 4 $10^5$[0.56] | Blood monocytes at baseline | This study (uninfected) |
| $k_{out_M}$ | /h | 0.04 | Elimination rate of blood monocytes | O'Connell et al. [S7] |
| $R_{in_M}$ | cells/mL/h | 1.6 $10^4$ | Production rate of blood monocytes | $M_{B_0} \times k_{out_M}$ |
| $M_{L_0}$ | cells/mL | 1.3 $10^6$ | Lung monocytes at baseline | Rodero et al. [S11] |
| $k_{M\,bl}$ | /h | 0.08 | Transfer rate of monocytes from blood to lung | This study* |
| $k_{M\,lb}$ | /h | 0.03 | Transfer rate of monocytes from lung to blood | $k_{M\,bl} \times M_{B_0}/M_{L_0} \times V_b/V_l$ |
| $M_{A_0}$ | cells/mL | 4 $10^6$[1.11] | Alveolar macrophages at baseline | Rodero et al. [S11], this study |
| $\alpha_{AM}$ | /h | 0.0015 | Renewal rate of macrophages | $k_{out_{AM}}/(1 - M_{A_0}/M_{max})$ |
| $f_{cM}$ | mL(lung)/mL(BAL) | 0.003 | Correction factor for macrophages | This study |
| $M_{max}$ | cells/mL | 1.2 $10^7$ | Macrophages plateau | Misharin et al. [S4] |
| $k_{out_{AM}}$ | /h | 0.001 | Elimination rate of macrophages | Murphy et al. [S6] |
| $E_{max,C}$ | - | 400 | Maximal effect of clodronate | This study (uninfected)* |
| $A_{50,C}$ | mg | 0.16 (34) [0.43] | Clodronate for half of maximal effect | This study (uninfected) |
| $d_M$ | /h | 0.6 | Transit differentiation rate of monocytes | Koogushi et al. [S3] |
| $K_{N_m}$ | CFUeq | 1 $10^5$ | Bacteria to reach half of max differentiation | Koogushi et al. [S3] |
| $f_{cB}$ | mL | 89 (23) [1.27] | Correction factor for bacteria | This study |
| $\alpha_S$ | /h | 0.75 | Maximal growth rate of susceptible bacteria | Roach et al. [S10] |
| $\alpha_R$ | /h | 0.75 | Maximal growth rate of resistant bacteria | This study* |
| $B_{max}$ | CFUeq | $10^{10}$ | Bacteria plateau | This study* |
| $\varepsilon_N$ | mL/(h.cell) | 2.9 $10^{-7}$(21)[0.75] | Neutrophil effect on bacterial decay | This study |
| $K_D, N$ | CFUeq | 5.4 $10^6$ | Bacteria to get half of maximal neutrophil effect | Roach et al. [S10] |
| $f_{cP}$ | mL | 1.5 (21) | Correction factor for phage | This study (phage decay) |
| $\phi$ | /(h.PFU$^{\gamma_P}$) | 1.2 $10^{-7}$(4)[0.07] | Adsorption rate of PAK P1 | This study |
| $\iota_M$ | - | 0.3 | Maximal adsorption inhibition by macrophages | This study* |
| $M_{50}$ | cells/mL | 2.4 $10^6$ | Macrophages to reach half of $\iota_M$ | This study* |
| $\gamma_M$ | - | 5 | Hill coefficient for adsorption inhibition | This study* |
| $P_C$ | PFU/mL | 7.5 $10^5$ | Phage adsorption saturation | This study* |
| $e$ | /h | 6 | Transfer rate between infected compartments | Previous one step growth study |
| $\beta$ | PFU/CFUeq | 100 | Burst size | Roach et al. [S10] |
| $\omega_P$ | /h | 0.023 (15) | Phage decay rate in lung | This study (phage decay) |
| $\psi$ | mL/(h.cell) | 1.1 $10^{-8}$(2) | AM-mediated decay rate of administered phages | This study (phage decay) |
| $\psi_n$ | mL/(h.cell) | 3.0 $10^{-6}$ | AM-mediated decay rate of produced phages | This study* |
| $\sigma_{inter_B}$ | $\log_{10} CFUeq/mL$ | 0.52 (9) | Error for bacterial density | This study |
| $\sigma_{inter_P}$ | $\log_{10} PFU/mL$ | 0.44 (28) | Error for phage density | This study |
| $\sigma_{slope_{MA}}$ | - | 0.50 (25) | Error for alveolar macrophages | This study |
| $\sigma_{slope_{NL}}$ | - | 0.44 (29) | Error for lung neutrophils | This study |
| $\sigma_{slope_{MB}}$ | - | 0.20 (20) | Error for blood monocytes | This study |
| $\sigma_{slope_{NB}}$ | - | 0.45 (15) | Error for blood neutrophils | This study |

RSE: relative standard errors (given in %), $\omega$: standard deviation of random effects (individual parameters followed log Normal distribution, see Methods), *: sensitivity analysis performed, AM: alveolar macrophage, $\sigma_{inter}$: additive error parameter, $\sigma_{slope}$: proportional error parameter.
